# The BeMoBIL Pipeline for automated analyses of multimodal mobile brain and body imaging data

**DOI:** 10.1101/2022.09.29.510051

**Authors:** M. Klug, S. Jeung, A. Wunderlich, L. Gehrke, J. Protzak, Z. Djebbara, A. Argubi-Wollesen, B. Wollesen, K. Gramann

## Abstract

Advancements in hardware technology and analysis methods allow more and more mobility in electroencephalography (EEG) experiments. Mobile Brain/Body Imaging (MoBI) studies may record various types of data such as motion or eye tracking in addition to neural activity. Although there are options available to analyze EEG data in a standardized way, they do not fully cover complex multimodal data from mobile experiments. We thus propose the BeMoBIL Pipeline, an easy-to-use pipeline in MATLAB that supports the time-synchronized handling of multimodal data. It is based on EEGLAB and fieldtrip and consists of automated functions for EEG preprocessing and subsequent source separation. It also provides functions for motion data processing and extraction of event markers from different data modalities, including the extraction of eye-movement and gait-related events from EEG using independent component analysis. The pipeline introduces a new robust method for region-of-interest-based group-level clustering of independent EEG components. Finally, the BeMoBIL Pipeline provides analytical visualizations at various processing steps, keeping the analysis transparent and allowing for quality checks of the resulting outcomes. All parameters and steps are documented within the data structure and can be fully replicated using the same scripts. This pipeline makes the processing and analysis of (mobile) EEG and body data more reliable and independent of the prior experience of the individual researchers, thus facilitating the use of EEG in general and MoBI in particular. It is an open-source project available for download at https://github.com/BeMoBIL/bemobil-pipeline which allows for community-driven adaptations in the future.

## Introduction

Electroencephalography (EEG) has rapidly evolved into a popular brain imaging method in various areas outside of traditional medical or psychological research. Recent developments in amplifier technology and improvements in data-driven analyses enable mobile EEG and Mobile Brain/Body Imaging studies (MoBI; Gramann et al., 2011, 2014), accelerating this progress. These methods do not only allow for participant movement (mobile EEG) but specifically address the interplay of active behavior and neural dynamics (MoBI) to better understand the neural foundation of embodied cognition (Jungnickel et al., 2019; Makeig et al., 2009). The electrical activity of the brain underlying cognitive processes is now investigated outside traditional laboratories in diverse scientific fields such as human factors (e.g. Protzak & Gramann, 2018), architecture (e.g. Djebbara et al., 2019), sports science (e.g. Büchel et al., 2021), clinical neuroscience (e.g. Short et al., 2020) and many more. As a result, researchers from various fields with manifold scientific and technical backgrounds handle data from multimodal brain-behavior assessments. This has led to a multitude of analytic approaches and analysis pipelines. In order to draw synergy from these exciting developments, we have identified the need for a standard pipeline that offers researchers the opportunity to focus on the actual research questions instead of overcoming methodological obstacles over and over again. To this end, we propose an easy-to-use and adaptive multimodal data processing pipeline, the BeMoBIL Pipeline, that enables documented, traceable, and objective data processing.

Despite some efforts to develop best practice guidelines (Chaumon et al., 2015) and to establish standard processing pipelines, especially for stationary recordings (Bigdely-Shamlo et al., 2015; da Cruz et al., 2018; Gabard-Durnam et al., 2018; Pedroni et al., 2019; Pernet et al., 2021; Rodrigues et al., 2021), a common basis in EEG data processing is still lacking (Robbins et al., 2020). In particular, pipelines targeting methodological obstacles such as data synchronization and subsequent processing of diverse data modalities (e.g. eye tracking data, motion capture) with different sampling rates are currently unavailable. In addition, more ecologically valid mobile EEG and MoBI protocols often come with increased noise levels in the recordings due to mechanical artifacts as well as biological activity stemming from the movement itself (Gramann et al., 2011, 2021; Richer et al., 2020). Handling increased noise levels in the recorded signal also requires new analysis approaches that often utilize information about the ongoing movement and thus rely on accurately synchronized multimodal recordings and specific analyses (Jungnickel & Gramann, 2016). While one toolbox for multimodal data analyses exists (MoBILAB; Ojeda et al., 2014), it is not supported anymore and lacks central analysis functions that are important for in-depth EEG processing and synchronized multimodal event extraction. Such data-driven event extraction, however, can be of central importance for highly realistic recordings. For example, eye blinks might be used for blink-based event-related analysis of EEG data when no external visual stimulation is available in natural outdoor experiments (Wascher et al., 2014; Wunderlich & Gramann, 2021).

This multitude of complex data types and the lack of common processing standards can lead to subjective, unjustified, or laboratory-specific parameter choices (e.g. filter design, artifact handling). As a consequence, peer-review processes can become complicated or, in the worst case, these factors can result in serious reproducibility issues (Cohen, 2017; Kappenman & Keil, 2017; Larson & Moser, 2017; Open Science Collaboration, 2015). We believe that standards in multimodal neuroscientific computing are the inevitable prerequisite for researchers to converge on basic principles in the field, further our understanding of human brain function, and foster more efficient research. The goal of the present paper is thus to introduce a replicable, open-access, standardized, and transparent analysis approach to EEG data in general and to multimodal mobile EEG/MoBI data analyses specifically. Our pipeline is intended to serve as an analysis basis that can be adopted and continuously developed by the scientific community.

The BeMoBIL Pipeline runs on MATLAB as we incorporated standard data processing routines from EEGLAB (Delorme et al., 2011; Delorme & Makeig, 2004) and Fieldtrip (Oostenveld et al., 2011), two of the currently leading EEG data processing toolboxes. Our pipeline can be used for the exclusive analyses of EEG data as well as for multimodal data processing including different data streams such as motion capture, eye tracking or force plate data. We provide raw data import and synchronization functions for the standard Brain Imaging Data Structure format (BIDS; Gorgolewski et al., 2016) that allows for easy data sharing. EEG data processing is available in the form of basic preprocessing routines (e.g. re-referencing, line noise removal, channel interpolation) as well as advanced artifact handling and source separation scripts (e.g. independent component analysis, ICA, or early-fusion approaches) and subsequent group-level source analysis methods. Motion data processing is integrated, including data preprocessing and the creation of derivatives. As a special feature for MoBI analyses, event extraction from motion data (e.g. heel strikes), eye tracking (e.g. blinks), electrocardiography (ECG, heartbeats), and EEG (using independent components representing eyes movements, heartbeats, or gait) are available. Various parameters can be adjusted for each processing step while the pipeline comes with informed recommendations for all steps. We encourage independent plausibility checks through automated data visualization at several processing milestones. All routines and parameter selections are clearly described and documented in a wiki on the GitHub repository (https://github.com/BeMoBIL/bemobil-pipeline). The source code is freely available and extensions and improvements from the community are welcome. Figure 1 depicts the general structure of the pipeline. In the following sections, each pipeline element is described in detail.

**Figure 1:**
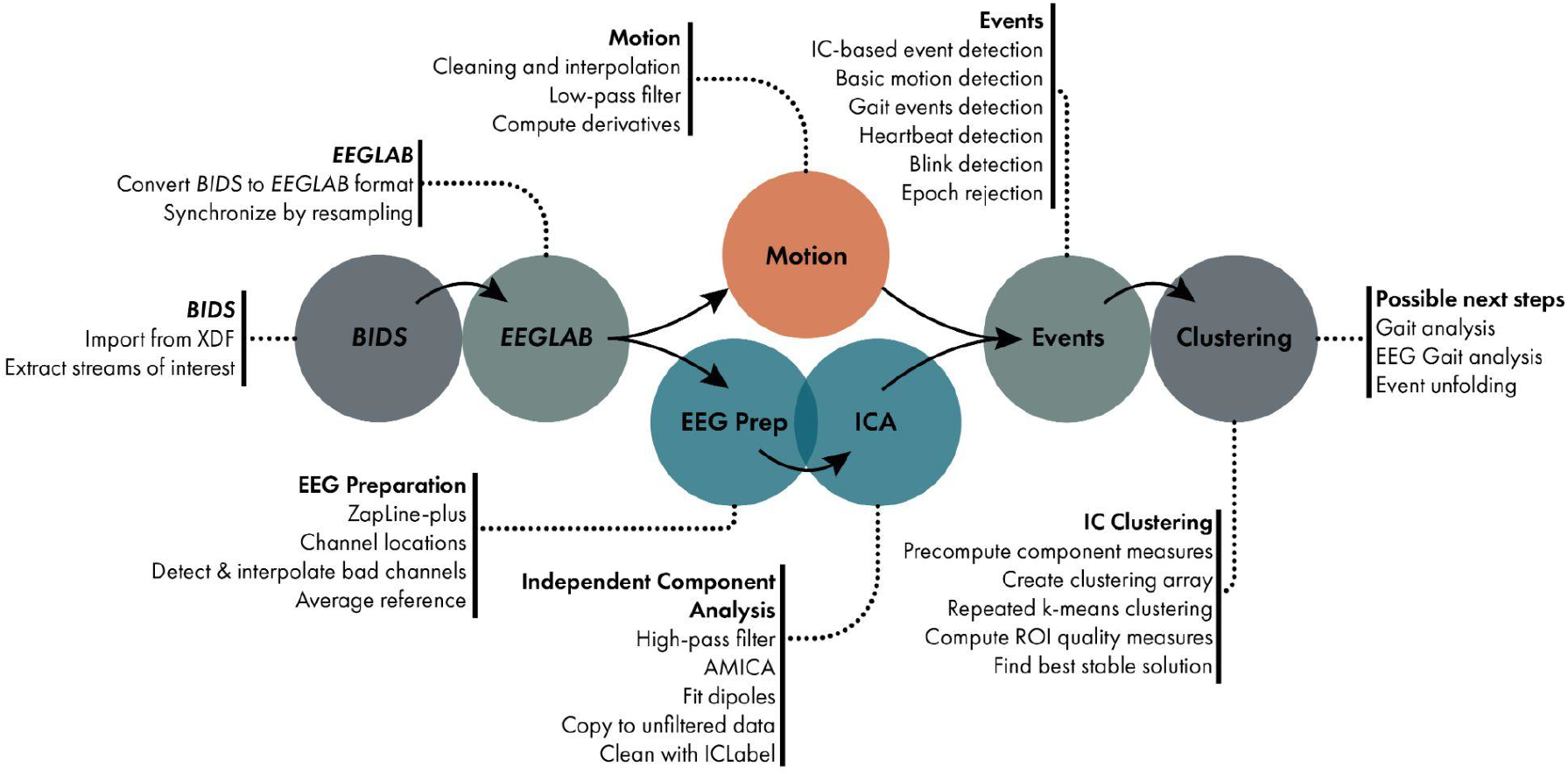
The BeMoBIL Pipeline workflow. Data is imported into BIDS format and then converted into synchronized EEGLAB files. EEG data is preprocessed, cleaned, and analyzed using ICA. Motion data is cleaned and prepared for analysis. Event markers can be extracted from different data modalities and repeated clustering can be used for robust source-level analysis. The individual steps are explained in detail in the respective sections.

## Time-synchronized data collection and import

Handling simultaneously recorded multimodal data requires special considerations from the stage of data collection and import. A harmonized, time-synchronized representation of EEG and motion data is a prerequisite for joint analysis of the two modalities. Addressing time synchrony is especially non-trivial in MoBI studies because a dataset often consists of multiple modalities of different sampling rates and different levels of time precision. For instance, although EEG recording systems typically yield highly regular inter-sample intervals, motion data may have irregular temporal distances between samples and may even completely miss latency information for each sample. One solution to record such multimodal data and preserve all available information is Lab Streaming Layer (LSL, available at https://github.com/sccn/labstreaminglayer). LSL is an open-source data streaming protocol that allows the flexible definition and recording of data streams from various sources such as different EEG amplifiers, different motion capture hardware, eye gaze recordings, experiment event markers, and more. Using the LSL recorder, the multimodal dataset can be stored in the extensible data format (XDF), containing all selected data streams with definitions, samples, and timestamps for each sample.

### Standardizing MoBI data

Although the XDF format is commonly used in the context of MoBI studies, it does not easily allow sharing the recorded data in a standardized way, as the recordings can contain self-defined data types with non-standard metadata and the XDF format is not based on common consensus. Additionally, the XDF files may not contain metadata about the experiment such as participant information that can be relevant for analysis. A key solution to easier data sharing is the BIDS standard. Initially covering standards for sharing fMRI datasets (Gorgolewski et al., 2016), the BIDS community has continued its effort to include more types of data in the framework by means of modality-specific extensions. The BeMoBIL Pipeline is designed to operate on datasets that adhere to the standards provided by the EEG-BIDS (Pernet et al., 2019) and MOTION-BIDS (in progress, see https://bids.neuroimaging.io/get_involved.html) extensions. Using BIDS data standards lowers the risk of data rot and makes data sharing easier by adhering to the FAIR principles for data management (Wilkinson et al. 2016), addressing the findability, interoperability, and reusability aspects. This in consequence can contribute to mitigating low reproducibility of scientific findings (Cohen, 2017; Kappenman & Keil, 2017; Open Science Collaboration, 2015).

To this end, the pipeline provides two features: One is the function *bemobil_xdf2bids* to convert data contained in XDF files into BIDS formatted data, and the other is the function *bemobil_bids2set* for converting BIDS data into EEGLAB-compatible data structures. The created BIDS dataset retains all necessary timing-relevant information from all streams, especially all timestamps of the samples, while applying only minimal change to the data itself, for example converting motion data orientation values that are represented in quaternion values into Euler angles, or replacing samples with a defined value for missing samples (e.g. zeros or -999) with not-a-number (NaN) values. The function supports the flexible use of parameters to load XDF files and allows the addition of metadata such as information about the participants, the EEG recording, the motion tracking system, its manufacturer, or the spatial axes layout to be stored together with the dataset.

When importing this BIDS dataset into EEGLAB using *bemobil_bids2set*, the function aligns the EEG and other data streams by first resampling the EEG data to a given target sampling frequency and then resampling other data to match the latency of EEG samples (see figure 2). As it can be assumed that the EEG data is recorded with high precision and equidistant samples, it is resampled to the desired sampling rate using the EEGLAB function *pop_resample*, which uses the filter-based *resample* MATLAB function internally. Resampling the other data types to the same rate is done by default using the *pchip* interpolation option of the fieldtrip function *ft_resampledata*, including an anti-aliasing filter if the data is downsampled. This is chosen as it is not always guaranteed that the nominal sampling rate of the other data types is accurate (e.g. when recording motion from virtual reality environments, the sampling rate is dependent on the performance of the rendering and the refresh rate of the display), and the samples are not always evenly distributed. Even in cases of equidistant sampling from reliable measurement devices, the level of precision using filter-based resampling may not be high enough for very long data sets containing millions of samples, leading to a shift between EEG and other data of several hundreds of milliseconds towards the end of the recording. In contrast, using interpolation of other data streams to align with the EEG samples preserves the relative temporal structure between the different modalities within the precision of one sample at all times and is thus favored in our use case. As motion data usually varies mainly in much lower frequencies than EEG, imprecisions introduced during the interpolation should not be problematic for downstream analysis. However, the requirements unique to additional modalities other than EEG and motion (such as eye tracking) may be at a disadvantage with this approach due to possible distortion of the signal, especially in the high-frequency range. In these cases, the exact sampling rate can optionally be entered and used for filter-based resampling instead of the interpolation approach.

**Figure 2:**
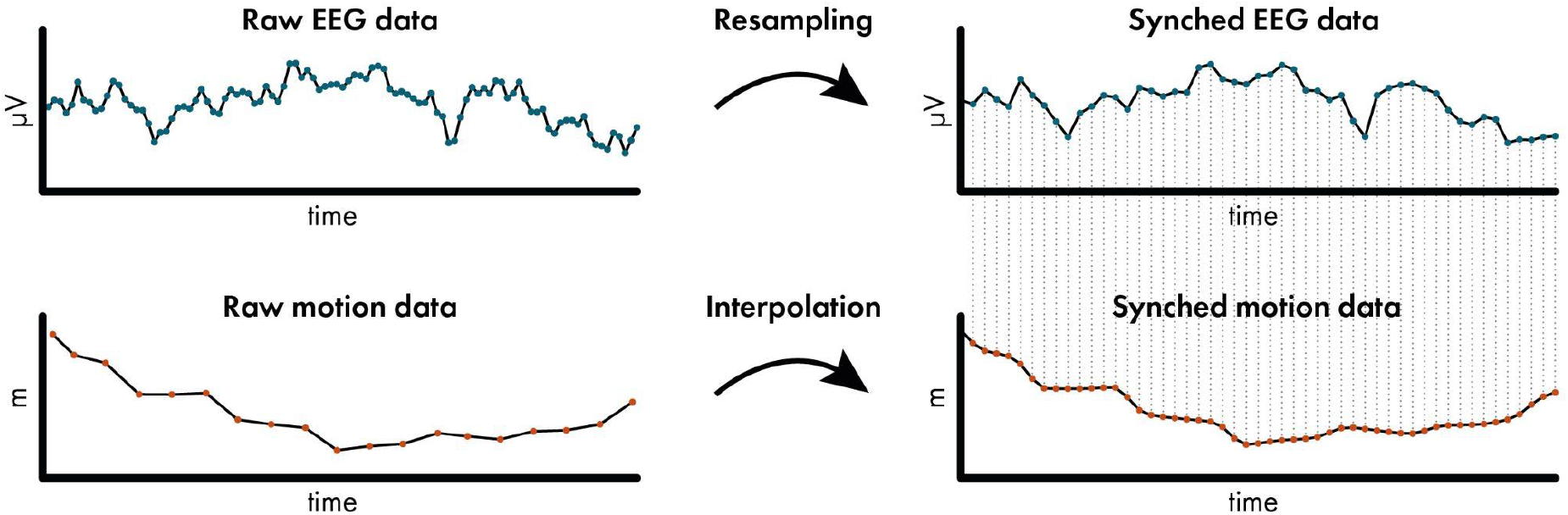
Example impression of the import synchronization. When importing BIDS data using *bemobil_bids2set*, EEG data is resampled using a filter-based method, while other data such as motion is interpolated to the same timestamps, leading to a fully synchronized dataset in EEGLAB. In this 200 ms example data, the EEG data was originally sampled at 500 Hz and the motion data at 90 Hz, and both are resampled/interpolated to 250 Hz. Plots of the raw and synchronized data that show the first channel of all modalities are created around the first and last event in each data file, allowing the inspection of the synchronization. This exemplary visualization is taken from subject 6 in the NeSitA example data that is available with the BeMoBIL Pipeline. For details, see section *Data import and time synchronization*.

For every recording then, a plot is created that shows the first and last event and one channel of all modalities, both directly imported from the XDF file as well as after the EEGLAB import and data alignment process. These plots can be used to verify the integrity of the temporal structure of the multiple data streams. As a final step of the import, all data modalities are made the exact same length as the EEG data, even in cases where there was no recording of a given modality in some sessions. To this end, all missing samples are filled with not-a-number (NaN) values. This, in combination with the previous resampling step, leads to fully synchronized data structures in EEGLAB with the exact same amount of samples and identical event markers in all modalities. Such synchronized data allows for analyzing event-related activity or the creation of event markers from one modality that can be copied to others, such as extracting gait event markers from motion (see section *Extracting gait parameters from motion data*). A generic template and a specific example script to use this complete import processing are available in the repository.

## Data cleaning and processing

The BeMoBIL Pipeline contains a variety of scripts and wrappers for cleaning and processing EEG, motion, and eye tracking data, all configured by one central file containing all relevant parameters and explanations thereof. EEG parameter default values are optimized for data from mobile experiments, however, they can also be easily adjusted for data from stationary experiments. Processing steps are documented within the data structures themselves, as well as by visualizing the outcome of every important step. As for the import, template files for the processing exist for all modalities and the configuration file. The pipeline wrapper scripts read the raw imported data and create several intermediate folders for preprocessing, other processing, and the final resulting data files. These folders and filenames can be adjusted in the configuration file. If the processing is stopped and restarted, it will by default load already created files instead of computing them again. This can be overridden if necessary. As it was shown that the data precision level has important effects on the processing (Bigdely-Shamlo et al., 2015), we ensure that double precision is used from the start and throughout the processing by selecting the appropriate EEGLAB option at several points within the pipeline.

### EEG data processing

The BeMoBIL Pipeline allows for fully automatic processing of EEG data from raw files to the final clean datasets including ICA information. This raw data can be obtained either via our own import (see above) or from any other importer. As a very first step in EEG processing, it is recommended, but not mandatory, to remove the segments before and after the experiment, as well as breaks during the experiment. This can be done automatically based on experiment event markers, or manually if no such event markers are included in the data structure. This step is important because these segments can contain very strong artifacts, e.g. excessive movement from stretching, or electrical or mechanical artifacts from touching cables, the cap, or putting off and on equipment such as a virtual reality head-mounted display. These artifacts can affect subsequent analysis steps negatively (such as channel rejection or ICA) and should thus be discarded. If removal based on event markers is impossible or manual removal is preferred for other reasons, it should be done once, and the removed indices should be stored for every participant. These indices can be obtained using the *eegh* command after removal, and can subsequently be applied automatically again. This is important on the one hand for reproducible processing, and, on the other hand, to be able to maintain synchronized datasets from different modalities. If segments are removed in the EEG data in this step, these segments must also be removed in motion and other physiological data for their respective processing. With only relevant experiment segments remaining in the EEG data, it is first preprocessed and then subjected to an ICA.

### EEG preprocessing

Preprocessing of EEG data in the BeMoBIL Pipeline is done using the *bemobil_process_all_preprocessing* function. This is a wrapper that incorporates all necessary processing steps from the raw loaded EEG set (all blocks merged together and non-experiment parts removed) up to the preprocessed dataset which has line noise removed, channels interpolated, and the data re-referenced to the average. It stores intermediate files on disk in the location provided in the configuration file parameter *bemobil_config*.*EEG_preprocessing_data_folder* and plots several analytics plots which are saved alongside their respective files. Exemplary visualizations of the preprocessing can be seen in figure 3.

**Figure 3:**
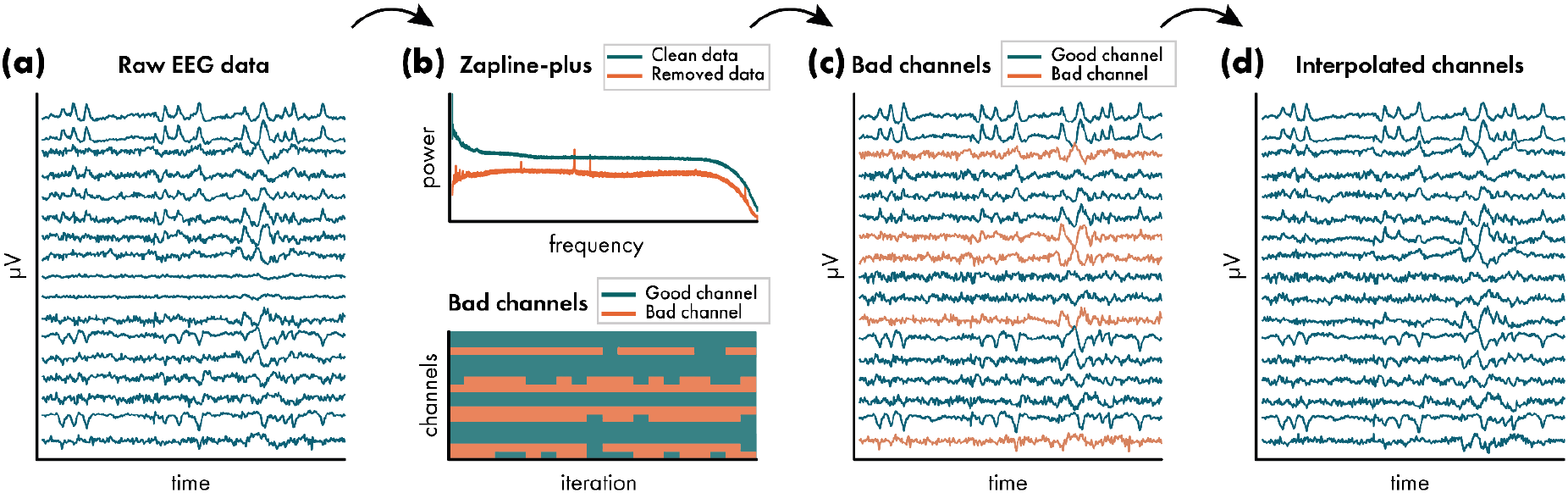
Exemplary impressions of the visualizations of the EEG preprocessing. During preprocessing of EEG data, several plots are created to allow an inspection of the workflow. (a) The raw data is plotted directly after import in six 10-second periods equally spaced throughout the entire data. This example shows the third such period in subject 76 of the visual discrimination datasets that are available with the BeMoBIL Pipeline. Only the first channels are shown in this plot for visualization purposes. Note that the scale of each channel is different and depends on the overall activity, and can thus not be interpreted. (b) Diagnostic plots of Zapline-plus (top) and the repeated bad channel detection (bottom) are available. This example shows a simplified version of plots from the same dataset as in (a). (c) In a plot similar to (a), the data that is cleaned with Zapline-plus is visualized with detected bad channels denoted in red color. (d) Lastly, the completely preprocessed data with interpolated bad channels is visualized again in a plot similar to (a). For details of the preprocessing, see section *EEG preprocessing*.

As a first step, some basic preparations are performed on the data using the *bemobil_process_EEG_basics* function. First, the EEG structure is filled with ur-data, specifically for event markers and channel locations. This is to ensure that in subsequent processing, the original event structure can always be recovered. Then, unused electrodes are removed. This can be set in the config file using the *bemobil_config*.*channels_to_remove* entry. This is to ensure that channels without channel location or channels in the montage that do not record data have no impact on downstream analysis. If the data is not already at the correct sampling rate, it is then resampled to the frequency declared in *bemobil_config*.*resample_freq*.

Frequency-specific noise is removed with Zapline-plus (Klug & Kloosterman, 2022). Zapline-plus is an EEGLAB plugin that removes spectral artifact peaks automatically. It includes a detector for artifactual peaks, chunks data to account for non-stationarity of the noise, runs Zapline (de Cheveigné, 2020) with the automatically detected number of components to remove, and creates a comprehensive analytics plot. Zapline removes noise by splitting the data into an originally clean (data A) and a noisy part (data B) by filtering the data once with a notch filter (A) and once with the inverse of said notch filter (B). It then uses a spatial filter to remove noise components in the noisy part (B) to get a cleaned version of that part (B’). Finally, the two clean parts (A and B’) are added back together to result in a cleaned dataset with full rank and full spectrum except for the noise. Zapline is preferred over a notch filter because it preserves the spectrum, and it is preferred over a simple spatial filter because it preserves the full data rank. It was also shown to have more powerful cleaning capabilities than the *cleanLine* plugin of EEGLAB (Miyakoshi et al., 2021). If *bemobil_config*.*zaplineConfig*.*noisefreqs* is declared empty, the function will use the complete automatic adaptation. This is the default and recommended. However, parameters can be adjusted if the cleaning does not work as intended. See (Klug & Kloosterman, 2022) for details about the processing and parameter tweaking. This step can be avoided by setting the whole *bemobil_config*.*zaplineConfig* field to empty.

After this first step of data cleaning, channel locations are added. Here, channel names can be changed in case they were named incorrectly or contain an unnecessary prefix using the *bemobil_config*.*rename_channels* setting. This is to ensure that the lookup tables for channel locations can operate correctly, even if the channel names were incorrect. At this point, a reference channel can also be added with zero entries when declared in *bemobil_config*.*ref_channel*. This allows feeding back the data of the reference channel when the data is re-referenced to the average in a later step, similarly to the *Full Rank Average Reference* EEGLAB plugin. This is done before importing channel locations so the reference channel can also be located. Then, if channel locations were not already loaded during the import, they can be imported at this stage using the *bemobil_config*.*channel_locations_filename* entry. In this step, we either look up locations in the standard 10-20 system (when no filename is provided) or use a file provided by an electrode location digitizer. In the latter case, if a reference channel was declared before, the file must contain the location of the reference with the name specified above. Lastly, the channel types are declared to be either EEG (default), EOG (can be provided in *bemobil_config*.*eog_channels* and will be ignored in both bad channel detection and re-referencing, see below), or REF (if a reference channel was entered above). These steps form the foundational preparations for downstream cleaning and analysis, and data files will be saved with the name provided in *bemobil_config*.*basic_prepared_filename*.

As the next step, bad channels will be detected and interpolated in a repeated process using the *clean raw data* plugin of EEGLAB, which uses the algorithm proposed in the PREP pipeline (Bigdely-Shamlo et al., 2015). We do not use all options in this function and our processing slightly differs from the PREP pipeline, which is why we make use of the general concepts only. One key issue in this process is the order of the re-referencing and interpolation, in which the average reference should not contain strong artifacts anymore, but re-referencing is necessary for the detection of bad channels. The detection of bad channels also recommends a high-pass filter of 0.5Hz cutoff frequency, which is not part of our preprocessing in order to preserve as much data information as possible, and since spectral filters should be adapted exactly to the needs of the final analysis (Widmann et al., 2015). For these reasons, we decided to split the detection of bad channels from their interpolation.

Using the *bemobil_detect_bad_channels* function, the data is first re-referenced to the average, excluding EOG channels as defined previously. This is to have an approximation of the final data but includes the impact of bad channels. This average reference will not be used later on, but only to detect the bad channels. Here, we either make use of the previously added reference channel or, as a fallback, the *Full Rank Average Reference* EEGLAB plugin, to preserve the full rank of the data and thus as much information as possible. This is automatically detected in the *bemobil_avref* function.

Subsequently, we repeatedly run the *clean_artifacts* function of the *clean raw data* EEGLAB plugin, with the number of repetitions specified by the *bemobil_config*.*chan_detect_num_iter* parameter. This is necessary because *clean_artifacts* uses a random sample consensus (RANSAC) approach that does not necessarily converge to the same results when repeated. The function stores the sampling in a micro cache that will be accessed when restarting the function without restarting MATLAB or clearing the micro cache beforehand, resembling a stable result. But when using the function with a cleared micro cache, the detected channels might differ. To ensure a reproducible bad channel detection, we thus clear the micro cache and repeat the detection several times, with a recommended minimum of 10 iterations. Only channels that were flagged as ‘bad’ more than a given proportion of the processed data (specified in *bemobil_config*.*chan_detected_fraction_threshold*) are then detected for final removal. We exclude all EOG channels from the detected bad channels because their statistical properties will often lead to false positive detection.

Within the *clean_artifacts* function, the data is split into windows of five seconds, and robust interpolations of each channel are computed based on the RANSAC sampling of surrounding channels. We do not make use of the time-domain sample removal or the Artifact Subspace Reconstruction (ASR) options, as we are only interested in detecting bad channels at this point. In our detection, five parameters can be adjusted:

- *bemobil_config*.*chancorr_crit* is the main parameter. This is a correlation threshold. If a channel is correlated less than this value to its own robust estimate based on the surrounding channels, it is considered abnormal in the given time window. Recommended are correlation values of 0.75 (rather lax) to 0.85 (rather strict).
- *bemobil_config*.*chan_max_broken_time* sets the maximum proportion of time windows a given channel may be flagged as bad before it is detected as bad in the final output per iteration. Recommended are values from 0.2 (20% of the time, strict) to 0.5 (50% of the time, lax).
- *bemobil_config*.*flatline_crit* uses a criterion for detecting channels that are flat. This is recommended to be set to ‘off’ since i) flat channels will not correlate with their interpolation, and ii) sometimes, especially in MoBI, data may be lost without necessitating the removal of the complete channel.
- *bemobil_config*.*line_noise_crit* rejects channels that have increased noise. However, line noise was removed by Zapline-plus in a previous step and it is recommended to be kept ‘off’, too.
- *bemobil_config*.*num_chan_rej_max_target* determines the fraction of channels that can be maximally removed (e.g. 1/5). This is to ensure that even in the case of very noisy data or incorrect bad channel detection, the processing does not remove too many channels to reconstruct them.

Subsequently, the detected bad channels are interpolated in the dataset that is not yet re-referenced or filtered using spherical spline interpolation in EEGLAB in the *bemobil_interp_avref* function. When this is done, the rank of the data matrix is reduced by the number of interpolated channels and this information is stored within the data structure. As a final step, the data is re-referenced to the average of all scalp channels (excluding EOG channels; Delorme et al., 2012), maintaining the full rank, as within the bad channel detection: Either the reference channel was declared previously, which means it was added with zero-entries. In this case, it will now be filled and available for analysis. Or no reference was declared, in this case, we follow the approach of the *Full Rank Average Reference* EEGLAB plugin: a new dummy channel with zeros is added, the data is re-referenced, then the dummy channel is deleted again. In both options, the data rank remains intact. The final preprocessed dataset is then saved with the filename provided in *bemobil_config*.*preprocessed_filename*.

### Independent component analysis

EEG measures not only brain signals, but always a mix of cortical, ocular, and muscular physiological sources, at times even cardiac activity in addition. Traditionally, any non-brain aspects of the data are considered artifacts and removed from the data, often in the time-domain. However, in MoBI data, physiological sources can carry important information and contribute to the interpretation of the data. Separating these sources and reconstructing their estimated activity throughout the experiment is thus an essential processing step, especially in mobile experiments where movement of the eyes and body is unrestricted. This can be done using blind source separation with ICA, which has become a staple in EEG analysis after having been introduced around three decades ago (Bell & Sejnowski, 1995; Hyvärinen & Oja, 1997). Different algorithms for ICA exist, but the Adaptive Mixture Independent Component Analysis (AMICA; Palmer et al., 2011) was shown to perform best in different comparisons, which is why we use it in this pipeline (Delorme et al., 2012; Leutheuser et al., 2013; Zakeri et al., 2014). The function *bemobil_process_all_AMICA* incorporates necessary steps from the preprocessed EEG dataset to the final dataset containing ICA information. The function stores the intermediate files of AMICA processing and accompanying plots in the folder specified in *bemobil_config*.*spatial_filters_folder* and its subfolder *bemobil_config*.*spatial_filters_folder_AMICA*. Exemplary visualizations of AMICA and subsequent processing can be seen in figure 4.

**Figure 4:**
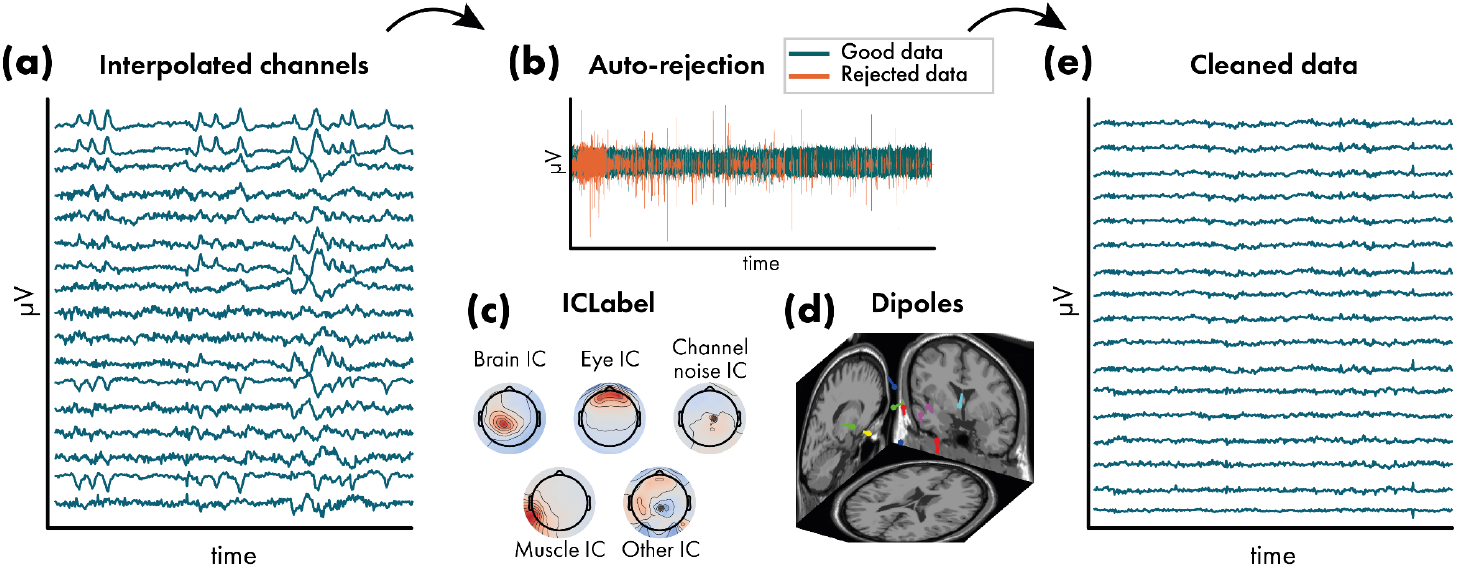
Exemplary impressions of the visualizations of the independent component analysis (ICA). During the computation of the ICA decomposition and subsequent source analysis, several plots are created to allow an inspection of the workflow. (a) The final preprocessed data is shown again here for visualization purposes (see Figure 3d). (b) An inspection plot of the automatic sample rejection of the AMICA algorithm is created (see section *Independent component analysis*), where rejected samples are denoted in red. (c) Exemplary visualization of the ICLabel classification of resultant independent components (ICs), see section *Original data reconstruction and independent component classification*. (d) Equivalent dipole models of the ICs that are classified as being of brain origin (subject 76 of the visual discrimination datasets that are available with the BeMoBIL Pipeline), see section *Equivalent dipole model reconstruction*. (e) Finally, the data is cleaned by removing unwanted ICs such as those stemming from non-brain sources, see section *Original data reconstruction and independent component classification*. The cleaned data is visualized in a plot similar to (a).

A vital step before computing ICA is to perform high-pass filtering with a suitable cutoff frequency. In earlier work, we showed that higher high-pass filters than commonly applied (> 0.5Hz cutoff) are often beneficial with diminishing returns, and that a higher cutoff frequency is recommended for data from mobile experiments and for higher channel density (Klug & Gramann, 2021). In addition, the improved decomposition also affects the signal-to-noise ratio of event-related potentials (ERPs) when cleaned with ICA, even when the ICA information was copied on an unfiltered dataset to make sure the effect was not due to the filter itself but only due to the improved ICA decomposition. Taken together, we found a filter of 1.25Hz cutoff to be a good overall option for data from both stationary and mobile experiments. However, for data containing strong movement, it might be suitable to use cutoff frequencies of 2Hz or even higher. It is important to note that the EEGLAB filter specifications do not use the cutoff frequency (the frequency where the filter starts taking noticeable effect, defined as -6db reduction in power) but instead, the user specifies the passband-edge (where the filter starts taking any effect). The transition bandwidth (two times the difference between passband edge and cutoff frequency) and resulting cutoff frequency are computed based on different heuristics depending on the frequency. It is recommended to specify the filter manually to ensure it leads to reproducible and comparable results (Widmann et al., 2015). We thus specify the filter order in the pipeline, and recommend using the same we used in our work comparing the high-pass filters (*bemobil_config*.*filter_AMICA_highPassOrder* parameter): The order of 1650 leads to a transition bandwidth of 0.5Hz, so to obtain a true cutoff frequency of 1.25Hz, a setting of 1.5Hz is required for the *bemobil_config*.*filter_lowCutoffFreqAMICA* parameter. A zero-phase Hamming window FIR filter is used. As with the high-pass filter for channel cleaning, this filter will only be applied to compute the AMICA and will not have any effect on the final dataset other than an improved decomposition. The final AMICA information will be copied to the unfiltered dataset.

Subsequently, the data is processed with AMICA. No time-domain cleaning is performed before doing so because AMICA includes a powerful cleaning option: At the beginning of the processing, samples with a log-likelihood estimation that is x standard deviations below the average (specified by *bemobil_config*.*AMICA_reject_sigma_threshold*, recommended 3 (Klug et al., 2022)) are removed repeatedly (as specified by *bemobil_config*.*AMICA_n_rej*, recommended 5 to 10 times (Klug et al., 2022)). This means that samples with suboptimal model fit are being rejected, as measured objectively by the algorithm itself. In our tests, this method yielded better or the same decomposition results than using our own time-domain cleaning option (see section *Time-domain cleaning based on epoch rejection* and supplementary material), which is why it is recommended and used in this pipeline. If desired, this feature can be disabled by setting the *bemobil_config*.*AMICA_autoreject* flag to 0. Note that these samples are not removed from the data itself, AMICA only disregards them internally when computing the spatial filters. A plot of the rejection is created and the rejection information is stored in the EEG data structure, thus being available for investigation or use in other circumstances. As it was shown, if stricter cleaning is being used (higher *bemobil_config*.*AMICA_n_rej* and/or lower *bemobil_config*.*AMICA_reject_sigma_threshold*), more samples will be discarded by AMICA (Klug et al., 2022). This can have a positive effect on the decomposition, which is important if subsequent source analysis is performed. However, in this case, the rejected samples will not contribute to the computation of the decomposition, which in turn means that the decomposition is not fit for these samples, and artifactual contributions might intersperse into brain components. Hence, it may be that even though the decomposition appears to be improved, the signal-to-noise ratio of the final measures (such as ERPs) is decreased or not changed in comparison to using more lax criteria when cleaning the data (Klug et al., 2022). We thus recommend using this time-domain cleaning conservatively: as much as necessary for the desired analysis, but as little as possible.

### Equivalent dipole model reconstruction

To obtain estimates of the source location for the resultant independent components (ICs), the fitting of equivalent dipole models is done using the *DIPFIT* toolbox of EEGLAB with standard settings of a 3-layer boundary element model (an exemplary visualization can be seen in figure 4d). If a non-standard electrode layout with individual electrode locations is used, these locations need to be warped to the standard locations to enable the correct fit of dipoles. For this, a subset of electrodes where the corresponding closest standard electrode is known can be entered in *bemobil_config*.*warping_channel_names*. The final dipole model has an accuracy of 1-2cm for brain ICs (Acar & Makeig, 2010) and includes information about the component topography variance that is not explained by the physiological model (residual variance, RV). The RV value can serve as an estimate of the physiological plausibility of the component and its respective dipole, and inform decision-making about the removal of components at a later stage (Delorme et al., 2012). Note that as we are interested in the location of not only brain but also eye and muscle source locations, it would be desirable to use a head model that does not restrict the sources to brain tissue only. This could be done using HArtMuT, a new head volume conduction model that extends to the neck and includes brain sources as well as sources representing eyes and muscles that can be modeled as single dipoles, symmetrical dipoles, and tripoles (Harmening et al., 2022). This model is currently under development and will be included in the pipeline as soon as it is available.

### Original data reconstruction and independent component classification

The ICA processing thus far was performed on high-pass filtered data. Since final EEG measures may require the data to be filtered with a lower filter cutoff (e.g. ERPs), the computed AMICA information including rejections and dipole fitting is copied back to the initial preprocessed, but unfiltered, dataset (Hyvärinen et al., 2001; Klug & Gramann, 2021). In order to provide a directly usable dataset, a final zero-phase Hamming window FIR filter can be applied using the *bemobil_config*.*final_filter_lower_edge* (high-pass filter, recommended 0.2 Hz) and *bemobil_config*.*final_filter_higher_edge* (low-pass, recommended empty, i.e. not used). This filter will be applied to all sets in the *bemobil_config*.*single_subject_analysis_folder*. Using the recommended 0.2 Hz high-pass filter will remove slow drifts in the data, but leave all relevant information intact, even for ERP analysis. Information about the filter is stored in the EEG data structure and can easily be reported. The filter order can optionally be specified, otherwise, EEGLAB default filters are being used. If both a lower and higher edge for the filter are entered, the filters are being applied successively without the use of a band-pass filter, as suggested by (Widmann et al., 2015). This dataset will then be stored in the folder specified by *bemobil_config*.*single_subject_analysis_folder* with the name specified by *bemobil_config*.*preprocessed_and_ICA_filename*.

As a last step in the EEG processing, the data is cleaned with the ICLabel classifier (Pion-Tonachini et al., 2019) as specified in *bemobil_config*.*iclabel_classifier*. ICLabel classifies ICs into brain, eye, muscle, and heart sources as well as channel and line noise artifacts and a category of other, unclear, sources, based on a large database of expert labelings (an exemplary visualization can be seen in figure 4). Our experience with MoBI datasets is that the ‘lite’ classifier detects muscle components more reliably than the ‘default’ classifier (Klug & Gramann, 2021). If it is only important to detect brain ICs, the ‘default’ classifier is likely to be the better choice. Only classes that are specified in *bemobil_config*.*iclabel_classes* are kept in the data, all others are removed. The majority vote is used by default, meaning each component is assigned the class with the highest probability. It is possible to also set a different threshold using the *bemobil_config*.*iclabel_threshold* parameter. In that case, the summed probability of the classes specified in *bemobil_config*.*iclabel_classes* must be higher than this threshold to keep the component. The cleaned dataset is saved with the name specified in *bemobil_config*.*single_subject_cleaned_ICA_filename* alongside a plot of the kept IC topographies and dipoles. Note that this is a critical step in the cleaning process. We found that ICLabel often performs sufficiently well to justify using it, since it makes the ICA cleaning objective and reproducible, but there might be cases where it fails, especially in datasets containing many muscle sources. A reason for this is that the original datasets used to train the ICLabel classifier were taken mostly from stationary experiments, thus muscle components or components related to mechanical artifacts stemming from movement are underrepresented in the classifier. It is very important that the resulting plots are inspected and checked for misclassifications. Guidelines for this process can be found in (Chaumon et al., 2015), and especially for MoBI data, a training tool for ICA labeling can be found at *https://www.icmobi.org*. On this website, experienced researchers can also label components in order to train a new classifier dedicated to MoBI data.

### Motion data processing

As for EEG, the BeMoBIL Pipeline provides a fully automatic processing pipeline for motion data from the raw data to cleaned and filtered data including derivatives. If motion data is to be analyzed in sync with EEG data with non-experimental segments removed, the same segments need to be removed in the motion data, too, to maintain the synchronization and enable easy transfer of event markers or other features between the two modalities. Data of rigid body movement can be processed using the *bemobil_process_all_motion* function. This function takes a full set of motion data, containing one or more rigid body tracked points with six degrees of freedom (3D position and orientation values, orientation can be given in quaternion units or Euler angles), and creates a cleaned motion dataset with orientation in Euler angles, containing derivative channels for velocity and acceleration in addition. The function performs the following actions:

1. Split the complete motion data into individual sets for each tracked rigid body, where each of the rigid bodies undergoes processing steps 2.-8.
2. Clean the motion data, which includes removing excessively large jumps in the data as well as interpolation of samples with lost tracking (NaN samples). This process also extrapolates data to parts of the experiments containing no motion information (NaN entries inserted during import, see section *Data import and time synchronization*) by entering the nearest available value.
3. Unwrapping Euler angles to eliminate jumps between -pi and pi in radian values. This is necessary for low-pass filtering, as otherwise, the jumps will create ringing artifacts.
4. Low-pass filter the data with the filter frequency given in *bemobil_config*.*lowpass_motion*.
5. Wrap the angles to pi again.
6. Compute the first derivative (velocity), ignoring the jumps from -pi to pi.
7. As time derivatives effectively amplify the high frequencies, it is recommended to use another low-pass filter after each derivative, as can be set in *bemobil_config*.*lowpass_motion_after_derivative*.
8. Compute the second time-derivative (acceleration) and add another low-pass filter as in 7.
9. Merge all single rigid bodies into one complete motion dataset again.
10. The final processed motion set is then stored on disk in the folder specified in *bemobil_config*.*motion_analysis_folder*.

## Event marker extraction

Experiments typically contain event markers to denote events within the experiment. This may be for example the beginning or end points of the experiment or experiment blocks, the presentation of stimuli, or responses of participants such as a button press. For stationary experiments in classical settings (seated participant, presentation of a stimulus on a computer monitor, no head movement), these event markers are sufficient for the investigation of most experimental questions. However, mobile EEG or MoBI studies inherently contain movement that may be explicitly part of the relevant measures. Here, extracting event markers post-hoc from the collected data can play an important role, as it can reveal information about cognitive and motor processes, or other physiological states. Hence, the pipeline provides a set of functions that allow easy and fast creation of event markers from the most prominent MoBI data types: Motion (general motion as well as gait events), eye gaze (blink events), and heartbeats. If the multimodal data was loaded via the pipeline import pathways, they will be completely synchronized, allowing extracting event markers in one data stream and copying them into another for analysis of event-related activity. Additionally, it is possible to generate event markers from the EEG data alone, even in the absence of other data streams.

### Motion event detection

Motion events can be an integral analysis aspect of MoBI experiments. For example, they can be used to detect reaction time and movement duration (Gehrke et al., 2022; Jungnickel & Gramann, 2016), they can serve as anchors for time-warping in spectral analysis (Gramann et al., 2021), they can help remove oscillatory gait artifacts (Gwin et al., 2010), or they can help to shed light on the neural basis of oscillatory gait generation (Wagner et al., 2016). To enable this functionality, the pipeline contains two functions: A basic movement onset and offset detector that requires only a single tracked element of any kind, and the advanced detection of relevant gait event markers.

### Basic motion event detection

The *bemobil_detect_motion_startstops* algorithm detects motion starts and stops based on a coarse and a fine threshold of one or more given channels. The square root of the sum of squares of these channels is taken as the detection data, resulting in the absolute value (if a single channel was entered), or the absolute movement in more than one dimension (if more than one channel was entered). An overall movement is detected first based on a coarse threshold of a given quantile of the data (0.65 by default), then a fine threshold is applied based on a buffer (plus and minus two seconds by default) around the detected initial coarse movement onset. This fine threshold is the minimum within the buffer plus a proportion of the range of the data within the buffer (0.05 by default). From the detected coarse movement onset going back in time, the last sample exceeding the fine threshold is taken as the final movement start, and from this point going forward in time, the last sample exceeding the fine threshold is taken as the movement stop. In effect, the coarse movement quantile threshold can be regarded as related to the amount of time one expects the tracked object to be in motion overall, while the fine movement threshold describes the expected data variability during the rest phase before the movement onset. The detector assumes no trend in the data and thus works on data where the endpoint of a movement is the same as the start point in the relevant channels. This can be for example the yaw orientation of the head to detect rotation movements (Gramann et al., 2021), the position of the hand in a reaching task (Gehrke et al., 2022), or the up/down movement of a foot tracker to detect steps (see next section). The detected event markers and used parameters are stored in the data structure so they can be copied between synchronized datasets of different modalities.

### Extracting gait parameters from motion data

Although computing final measures of specific movements is not in the scope of the BeMoBIL Pipeline, we would like to point out a few standard parameters that are commonly investigated in a number of mobile EEG and MoBI experiments. The aim of including this description is to enable researchers that are familiar with EEG, but not motion analysis, to obtain results comparable to biomechanics research. A large number of previous mobile EEG and MoBI studies investigated human brain dynamics in walking participants demonstrating the importance of gait in the research field. Here, one branch investigates the cortical dynamics accompanying gait control (e.g. Castermans et al., 2014; Gwin et al., 2010; Seeber et al., 2015; Wagner et al., 2012). A second branch of studies rather uses gait to investigate human brain dynamics in ecologically valid scenarios (e.g. De Sanctis et al., 2020; Jacobsen et al., 2021; Malcolm et al., 2015; Nenna et al., 2021; Protzak et al., 2021; Protzak & Gramann, 2021; Reiser et al., 2019, 2021). Here, the focus is less on gait itself but rather on the extraction of gait parameters in different walking scenarios to clean the signal from gait-related artifacts or to investigate the impact of walking difficulty on brain dynamics, respectively. Both approaches, however, require reliable extraction of gait parameters to inform the EEG analyses. To facilitate the extraction of reliable parameters and to provide guidelines for future mobile brain imaging studies, the definitions of a number of biomechanical gait parameters are given here.

As the main categorization, gait parameters could be divided into parameters of pace (including gait velocity or walking speed and referring step length), rhythm (including cadence, stride, and swing time), phase (including double support time), base of support (including step width) as well as variability (including the coefficients of variation for all parameters; Hollman et al., 2011). From a biomechanical point of view, an active heel-to-toe movement with associated ankle movement is necessary to maintain balance during the forward motion of walking. At the same time, with each step, the body’s center of gravity shifts beyond the support surface, so that the pelvis must be stabilized in the period from heel strike to double stance phase (Perry & Burnfield, 2010). Therefore, the movement of the feet, as well as the pelvis and hip rotation, give insights into stable gait patterns. Nevertheless, gait is a whole-body movement, and therefore trunk rotation, head movements, and arm swing and their referring kinematics are oftentimes relevant aspects to consider, e.g. to detect pathological gait patterns.

To describe the susceptibility to disturbances of gait and its accompanying brain dynamics, a set of parameters has been established quantifying changes in various gait parameters. These include i) reduced stride length, defined as the distance that one part of the foot travels in front of the same part of the other foot during each step, we recommend using the distance from heel-strike to heel-strike (Hollman et al., 2011; Scott et al., 2015), ii) reduced walking speed (Verghese et al., 2009), iii) prolonged stance phases, e.g. expressed by the double support time (the time when both feet are in contact with the ground simultaneously), defined as the sum of the time elapsed during two periods of double support in the gait cycle (Hollman et al., 2011; Maki, 1997; Scott et al., 2015; Verghese et al., 2009), and iv) increased stride length variability, defined as the coefficient of variation (%CV), calculated as the average standard deviation of the gait parameter divided by the average mean (Hollman et al., 2011; Verghese et al., 2009).

An essential requirement for measuring and calculating spatiotemporal gait parameters (i.e step length and cadence) is the accurate spatiotemporal identification of heel-strike and toe-off events (Rudisch et al., 2021). We provide options to extract these event markers, but no additional gait parameters, as these can be derived from the data and event markers but may require knowledge about the measurement device or experimental paradigm that is impossible to anticipate in a generalized pipeline. Due to underlying differences in measurement devices and principles, it has to be ensured that the gait cycles can be accurately detected by using a standardized description of the axes in the Euclidean space. In biomechanical analyses, the gait parameters are commonly defined such that the x-axis describes the anterior-posterior direction, the y-axis describes the medial-lateral direction, and the z-axis refers to the vertical direction (distal and proximal, up and down). With these axes known, gait event markers can be extracted in a standardized way using the *bemobil_gait_analysis* function.

The BeMoBIL Pipeline extracts gait cycles defined as a sequence of events: 1) movement start (flat foot phase end), 2) toe-off, 3) heel-strike, and 4) movement stop (next flat foot phase start). For an ideal toe-off and heel-strike event detection, the heels and toes would require their own tracker. However, these events can be reasonably approximated by assuming that i) during the flat-foot phase the foot is moving backward in relation to the body, ii) as soon as the toe-off event occurs, the foot starts moving forward, and iii) as soon as a heel-strike event occurs, the foot stands still again, and is moving backward in relation to the body. Detecting these four events thus is possible with only one tracker on top of the feet and happens in two steps: First, foot movements, in general, are detected using the *bemobil_detect_motion_startstops* function on the z-Axis of the tracking (up-down movement). These mark the flat-foot end and the flat-foot start events, respectively. In a second step, toe-off and heel-strike events are then defined using the velocity in the x-Axis (forward-backward). This requires the foot movement measurement to be in relation to the body, i.e. not a continuous forward movement but a forward-and-back cycle. If the motion was measured on a treadmill, this is already the case (as the feet slide back under the body). In overground walking, a motion tracking of the torso or head of the participant is required in addition. If such tracking is provided, the values in the x- and y-axes are subtracted from those of the feet, such that the feet exhibit a forward-and-back cycle again, relative to the provided tracking. With this cyclic movement in the x- and y-direction, one additional issue has to be overcome: The tracking axes are not necessarily always aligned with the movement axes, but the movement x-axis is relevant for event extraction. Hence, a PCA analysis is computed using the provided x- and y-axes for each foot separately, and finally, the component with the higher variance is taken as the foot x-axis. In this oscillatory forward-backward movement of the feet, the zero-crossings of the first derivative (i.e. maxima/minima) are taken as the final two events: Such a zero-crossing after the foot movement start event is used as the toe-off event, and the same before the foot movement end event is used as the heel-strike. The final gait event markers are added to the EEGLAB data structure and can be copied to synchronized other datasets such as EEG, allowing further investigations. An example visualization of this gait event extraction can be seen in figure 5.

**Figure 5:**
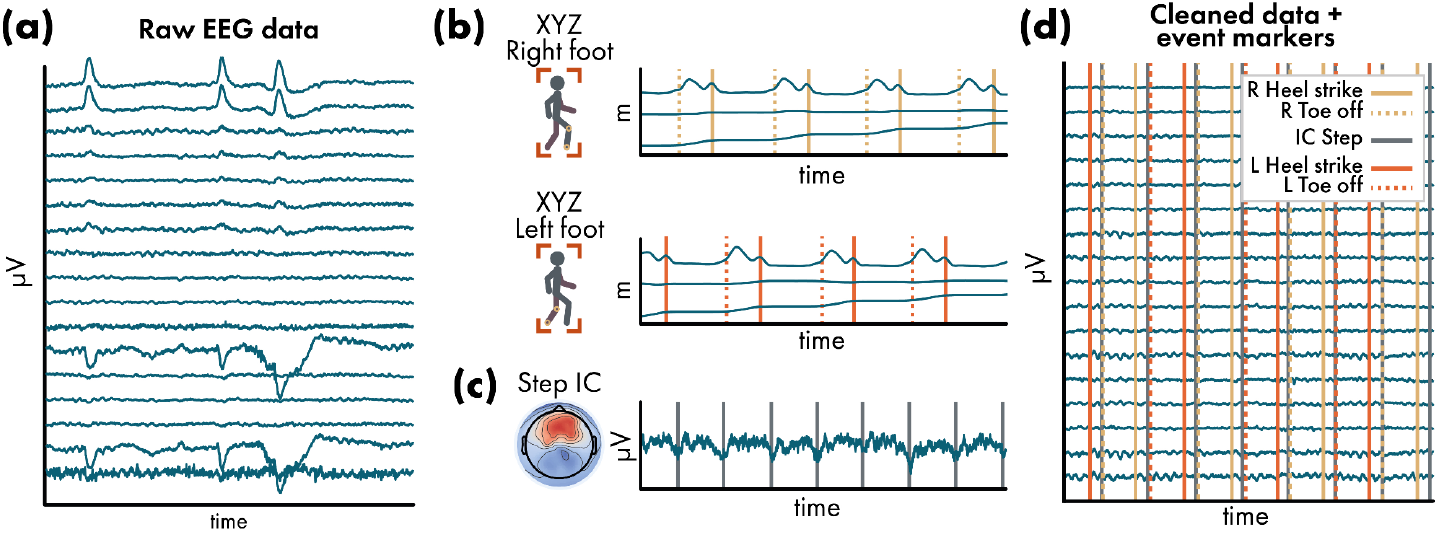
Exemplary impressions of the motion event extraction. (a) Example visualization of a 5-second data segment of raw data from subject 64 of the visual discrimination datasets that are available with the BeMoBIL Pipeline. (b) Motion data with gait event extraction (here, only toe-off and heel-strike events are plotted for visualization purposes). The x-axis of the feet is at the bottom of the two plots, respectively, and shows the forward movement. The y-axis represents the left-right movement of the foot, which is minimal in this case. The z-axis at the top of the two plots shows the lifting of the feet, with a lowering during the swing phase and a prolonged minimum during the flat foot phase. For details on the extraction of these events, see section *Extracting gait parameters from motion data*. (c) Exemplary detection of steps of both feet based on the independent component time series, see section *IC-based gait-related event detection*. (d) Exemplary final cleaned EEG data similar to (a) that contains all gait events for further analysis. Since the data modalities are synchronized during import (see section *Data import and time synchronization*), the events can easily be transferred between them.

### Eye tracking-based blink event detection

Eye gaze data can be very informative for MoBI analysis, and especially using VR displays it is easy to be recorded. During blinks, the pupil is not detectable, which means that the eye tracker can not determine the gaze direction. For a basic eye gaze analysis, the *bemobil_clean_eye* function thus cleans eye gaze data by detecting blinks based on the pupil radius. It then interpolates all eye gaze data using *pchip* interpolation during the detected blink times. Eyeblink event markers are added to the event field of the dataset and blink extraction information is stored.

The blink detection is modified from code written by Ravi Chacko (Mitz et al., 2017) and processes the data as follows:

1. The pupil radius is used to determine blinks, as during closure of the eyelids the pupil radius is zero. To this end, the mean and SD of the radius throughout the recording are computed, disregarding samples below a radius of 0.2mm. The general threshold to define a blink is then defined in SDs below the mean, 3 by default.
2. Going through the data from the start, a blink is detected if a sample is below the threshold. The following first sample above the threshold is regarded as the end of the blink.
3. These coarse start and end values are now used to determine the exact starts and ends. Here, a search buffer around the detected timestamps is used (20ms by default). Within this period, large peaks in the absolute values of the first derivative of the pupil size are computed and the last peak denotes the true end of the blink. This allows for a brief period in which the pupil size can jitter after re-opening the eyes (e.g. several samples in which the pupil radius appears to be open, then closed again, then open again), but only the final opening is taken as the end of the blink.
4. A buffer around the detected start and end points of the blink is applied (30ms by default), the blink indices are stored, and the detector continues after the end index with 2.
5. After searching for blink indices in the entire dataset, the detected blink periods are interpolated using *pchip* interpolation in all eye tracking data. This also removes large jumps in the eye gaze position data that can occur during blink periods, when the pupil is not trackable.
6. Finally, blinks that were below a minimum duration (100ms by default) are discarded, and the final blink start and end event markers are added to the EEGLAB data structure. These can be copied to synchronized other datasets such as EEG, allowing further investigations. An example visualization of this blink event extraction can be seen in figure 6.

### Event detection based on independent component time series

Paradigms including mobile EEG/MoBI can be comparably complex and time-consuming so it can be reasonable to reduce the data recording to EEG only. Even though this limits the extraction of event markers based on other modalities such as motion or eye gaze, there is another possible option to extract event markers for further analysis utilizing the decomposition of the data using ICA. Although ICA is commonly applied to remove non-brain activity from the data, it can also be used to extract event markers from components that stem from eyes, muscles, or mechanical artifacts like cable sway or electrode pressure from walking. Hence, these non-brain sources can now inform the analysis of brain activity by providing context events. Thus what is traditionally considered an artifact can become a signal and thus an integral part of the data analysis.

**Figure 6:**
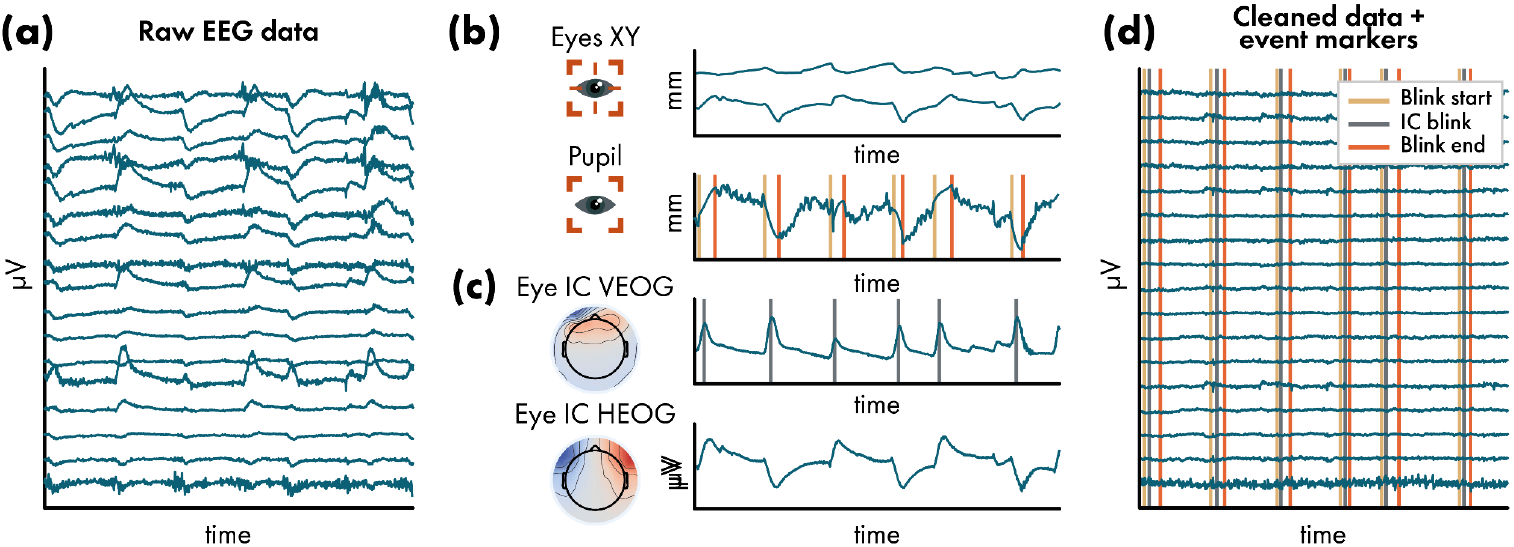
Exemplary impressions of the blink event extraction. (a) Example visualization of a 5-second data segment of raw data from subject 6 in the NeSitA example data that is available with the BeMoBIL Pipeline. (b) Eye tracking data with blink event extraction. Blinks are detected based on pupil size (see section *Eye tracking-based blink event detection*). The eye x- and y-axes are not used for event extraction but are shown for the visualization of the synchrony of eye tracking and independent component (IC) time series in (c). (c) Exemplary detection of blinks based on the IC time series, see section *IC-based blink and saccade event detection*. (d) Exemplary final cleaned EEG data similar to (a) that contains all blink events for further analysis. Since the data modalities are synchronized during import (see section *Data import and time synchronization*), the events can easily be transferred between them.

### IC-based blink and saccade event detection

Following the approach published in Wunderlich & Gramann (2021), the BeMoBIL Pipeline provides two functions to extract blink, saccade, and step event markers from IC time series. The function *bemobil_detect_blinks_from_ICA* detects blinks and saccades based on the time series of two ICs representing horizontal and vertical eye movements, respectively (e.g., figure 6c). In the first step, these components are automatically detected based on their topographies and their spectral power below 5Hz. It is also possible to indicate specific ICs manually and hand them over as parameters when using the function. Blinks and saccades are detected using the *findpeaks* function. Default values for distance, prominence, and width of the peaks are provided based on EOG literature (Lins et al., 1993). Alternatively, the parameters can be set by the user. Before detecting peaks in the IC activation time course, the data is smoothed using a moving median filter, which preserves the steep edges while removing high frequency fluctuations (Bulling et al., 2011). The moving median filter *smoothdata* is used with a window length of 0.08s by default and can be defined by the user. Furthermore, *bemobil_detect_blinks_from_ICA* takes care of flipped IC activity and ensures that blinks are always positive peaks which is a requirement for the use of the *findpeaks* function. For saccade detection, the square root of the sum of vertical and horizontal eye movement is computed which is known in the electrooculogram (EOG) literature as EOG activity (Jia & Tyler, 2019). The squared derivative of the EOG time series allows for using peak detection to locate the quickest differences in the EOG time series equalling saccadic eye movements. To disentangle the blinks from saccades, all saccade detections in temporal proximity (by default +/- 100ms) of a blink are excluded. For all the remaining detected peak latencies, EEGLAB event markers are created using the type ‘blink’ or ‘saccade’, respectively.

An informed decision about the detector efficacy can be made by the provided figures. Here, one plot shows the activation of the two detected eye ICs and the newly created event markers. In addition, there are figures for blinks and saccades, respectively, depicting the whole dataset with the *findpeaks* parameters, allowing the inspection of the peak detection efficacy when zooming in. Below this plot are histograms of the prominences and peak widths (including those exceeding the thresholds). These histograms provide information about how well the used threshold fits this participant’s data. In our tests and comparisons with eye tracking data in an experiment containing strong eye movements, we found the IC-based blink detector to be in correspondence with the eye tracking-based detector (see section *Eye tracking-based blink event detection*) around 80% of the time. This, however, depends on the nature of the eye movement, as e.g. strong vertical movements can appear almost like an eye blink in the vertical IC activity even in the absence of a blink. Saccades can thus falsely be detected as blinks, mask blinks, or go undetected because they can happen during a blink. Taken together, the event extraction should always be handled with care and especially saccades are not always reliable, which is why we offer an option to not extract saccade event markers. All parameters, the detected blink and saccade event latencies, as well as the prominences and widths of the detector, are stored in the EEG data structure. An example visualization of this IC-based blink detection can be seen in figure 6.

### IC-based gait-related event detection

The function *bemobil_detect_steps_from_ICA* detects steps based on the time series of an indicated gait IC (e.g., figure 5c). The most indicative signature of a gait IC is that the time series follows the same pattern as the upwards axis of a motion tracker device mounted to the head. As some data might contain steps only in parts of the entire duration, it is possible to specify the start and end points of the step detection to prevent false alarms during periods where the participant did not walk. Analogous to the eye-movement detector, the data is smoothed using the moving median and the function checks whether the stronger deflection is plotted upward. Steps are detected using *findpeaks* with parameters for distance, prominence, and minimal and maximal duration of the peaks. Finally, EEGLAB event markers are created at the respective latencies and stored in the EEG data. The parameters can be chosen freely and an informed decision about the detector efficacy can be made by the plots of the detection, including a histogram of the prominences and widths (including those exceeding the thresholds) akin to that of the blink detector. The detection can be repeated with varying search boundaries or IC indices. All parameters, the found step event markers, and latencies, as well as the prominences and widths of the detector, are stored in the EEG data structure. An example visualization of this IC-based step detection can be seen in figure 5.

### Heartbeat event detection

Heartbeats and subsequent analyses such as heart rate variability can be of interest in a MoBI experiment, for example for assessments of workload and stress (Delliaux et al., 2019; Kim et al., 2018). We thus provide the widely used Pan-Tompkins algorithm to detect heartbeats from electrocardiography (ECG) data (Pan & Tompkins, 1985), either recorded from additional ECG sensors or derived from independent components reflecting cardiac activity. As the original algorithm is written in the C programming language, we use a modified version of a MATLAB implementation available online (Sedghamiz, 2014), additionally allowing the specification of the high and low-pass filter cutoff frequencies (1 Hz and 40 Hz by default, respectively). The MATLAB implementation by Sedghamiz (2014) makes use of the hard-coded frequency-dependent filters and computations defined in the original work if the data is given at a sampling rate of 200 Hz, but uses state-of-the-art MATLAB signal processing otherwise, which is our recommendation.

## Single-subject and group-level post-processing

The BeMoBIL Pipeline focuses on the automatic processing and cleaning of EEG and other data but provides a selection of useful additional features regarding the next steps. With preprocessed EEG data as well as event markers available, this often is to analyze the data based on epochs that are centered around one or several events of interest. Using these epochs, it is then possible to perform analyses on either the cleaned sensor data or on the source-level, taking into account the location of the EEG equivalent dipole models of the ICs. To facilitate these steps, two options provided by the pipeline are helpful: First, we offer a way to reject epochs based on objective criteria and in a balanced fashion between conditions, and second, the pipeline includes a repeated IC clustering approach for reliable and reproducible group-level source analysis.

### Epoch rejection and time-domain cleaning

After creating epochs either based on experiment event markers or based on the event markers created using the pipeline, these epochs might still contain non-brain signals even if the data was cleaned with ICA before. As a final option to improve the signal strength of the measure of interest, it is thus often necessary to reject epochs that are particularly noisy. This can be achieved by either manually selecting epochs to reject, or by using automated methods. One issue arising in many automated rejection tools, however, is that one cannot specify the amount of data to be removed, but only the threshold that leads to removal. Hence, one runs the risk of insufficient cleaning or the removal of an excessive amount of data when the threshold is not adjusted properly. More importantly, cleaning data from different movement conditions might lead to an imbalance in the removal of epochs, where significantly more epochs are rejected in the condition with more movement, potentially complicating the analysis or skewing the final results.

#### Automatic and balanced epoch rejection

We thus propose a method to rank epochs on their noise level and remove only a specified amount of the worst ranking epochs. Each epoch is evaluated using four measures that are normalized by their median across epochs: i) the mean of channel means, to catch epochs with high amplitude, ii) the SD of means, to catch epochs with inhomogeneous channel activity according to their mean, iii) the mean of SDs, to catch epochs with high variance within channels (e.g. strong leftover muscle activity), and iv) the SD of SDs, to catch epochs with inhomogeneous channel activity according to their variance. It is possible to weigh the measures separately, although the default of equal weights is recommended. Each epoch then receives a final summed score and the epochs are sorted according to that score. Then three options are available to determine the rejection threshold: i) a fixed number of epochs that should be left - this will guarantee an equal number of epochs for all conditions, ii) a fixed percentage threshold, e.g. the worst 10% of the epochs are removed - this will preserve the original ratio of epochs per condition, or iii) determine a “knee-point” of the score and use that as the threshold - this will lead to the removal of only outlier epochs. Downsides of the third method are that this can lead to an imbalance of the retained epochs between conditions, and in cases where very few very strong outliers exist, the “knee-point” can be shifted to a high threshold, while very clean datasets can exhibit an almost round curve with a “knee-point” that is shifted towards the center. Thus, we recommend using methods i) or ii).

#### Time-domain cleaning based on epoch rejection

The algorithm to reject epochs can be extended for use as general time-domain cleaning of continuous data. To this end, the data is first high-pass filtered and subsequently cut into epochs that are then cleaned as described above. If eye movements are to be ignored in this cleaning, it is recommended to use a high-pass filter of 10 Hz to remove the majority of eye contributions. This, however, is unnecessary if the cleaning is used on data where eye contributions were removed with ICA. To target bad segments more precisely, epochs can be specified to overlap such that a short burst of noise could be captured by one epoch rather than two adjacent ones. Additionally, when an epoch is marked for rejection, a buffer around the epoch is rejected as well to capture possible on- and offsets of the artifact. The epoch length, overlap, and buffer can be specified according to the needs of the analysis. For example, when removing artifacts before running ICA, an epoch length of 500 ms with an overlap of 125 ms and an epoch buffer of 62.5 ms can be used, while longer epochs could be useful if it is important to retain longer contiguous data. We do not recommend using this method in our pipeline because we found no improvement when using it over the automated cleaning of AMICA itself.

### Robust group-level source analysis for regions of interest

If source analysis is to be performed on the group level, it is necessary to find ICs of all participants that represent activity from the specific source region. To this end, k-means clustering can be used, which finds similar components in the complete study set containing ICs from all participants based on weighted measures such as dipole location, scalp topographies, spectrum, ERPs, or event-related spectral perturbations (ERSPs). Choosing the weights is subject to the analyst, but it is recommended to weigh the location highly, add topographies and spectra, and, depending on the situation, ERPs and ERSPs with lower weighting. However, when using ERPs or ERSPs, it can be argued that double-dipping happens in the selection of relevant ICs (meaning that the measure that is later used to compute statistics is also used to select the ICs). A counter-argument to this would be that the clustering uses average measures while the statistics are used to investigate condition differences. All in all, no final rule on how to choose the weights can be given. The standard k-means clustering, however, has one other strong limitation: the k-means results are not stable due to variation in the starting conditions. Repeating the clustering can result in different solutions, and depending on the location and the similarity of the ICs, the cluster of interest (COI, the cluster closest to your region of interest, ROI) can contain vastly different ICs (see figure 7a).

**Figure 7.**
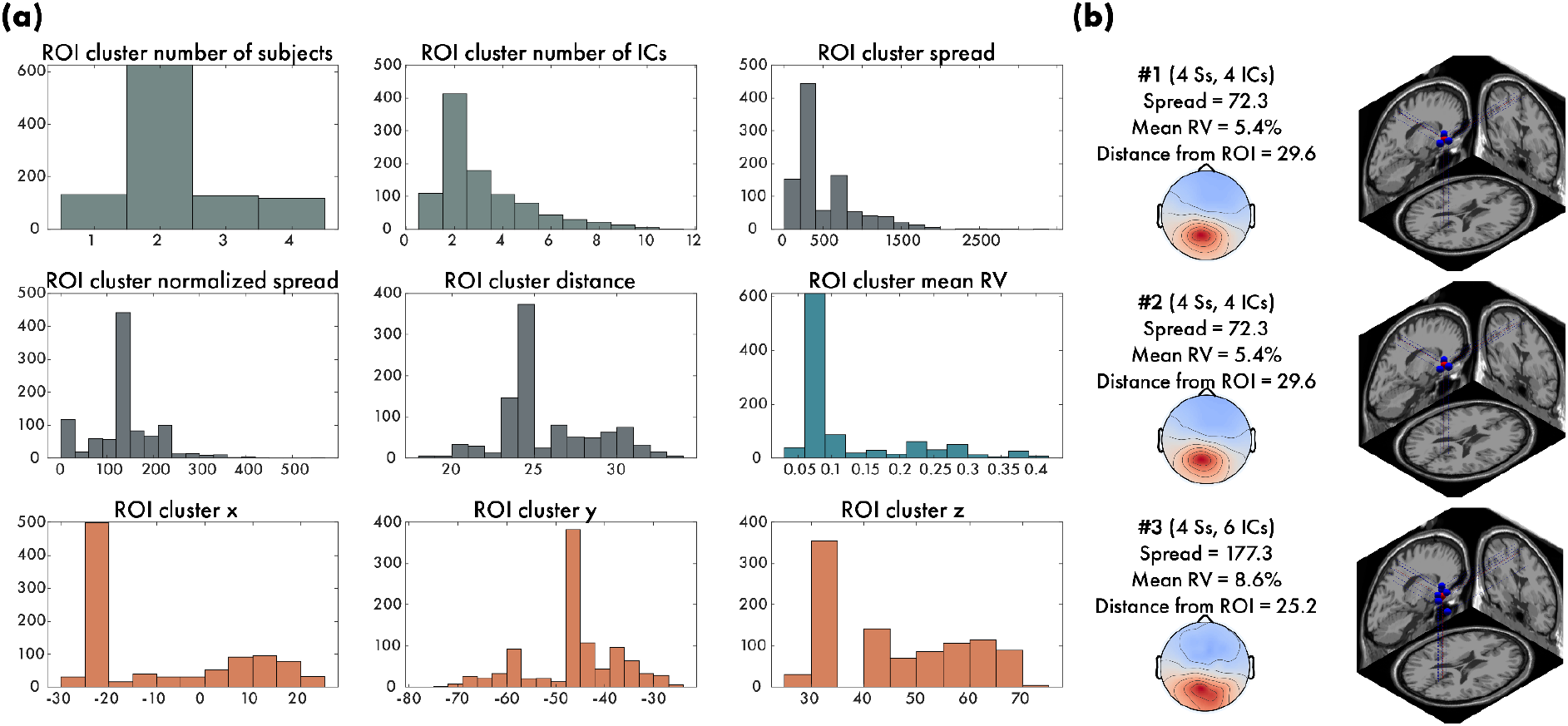
Robust group-level clustering example. In this example, we ran the robust clustering of independent components (ICs) on the group level of the four example participants from the visual discrimination study that is available with the BeMoBIL Pipeline. We weighted the dipoles with 3, the topographies with 1, the spectra with 1, and the ERPs of stimulus presentation with 1, and ran 1000 repetitions of the clustering. The region of interest (ROI) was set to the posterior parietal cortex (MNI coordinates of [0,-48,39]) and the weights for the ROI cluster quality measures were chosen as: subjects = 3, ICs/subjects = -1, normalized spread = -1, mean RV = -1, distance from ROI = -2, and mahalanobis distance from median of multivariate distribution = -1. See section *Robust group-level source analysis for regions of interest* for details. (a) The resultant distributions of the various quality measures show the variability of the outcome of the clustering. The unnormalized spread and the X/Y/Z coordinates are not used for the final selection but shown for visualization purposes only. (b) The resulting best three clusters and their respective measures show that, although they are not identical, they are similar, and the top 2 clusters are identical, indicating a stable result of the repeated clustering.

To alleviate this issue, we implemented a repeated clustering approach that clusters several hundred or thousand times and selects the COI for each clustering solution, based on the distance from a given ROI in MNI coordinates (Evans et al., 1993). For each of these COIs, a set of quality measures is derived: the number of subjects in that cluster, the average number of ICs per subject, the normalized Euclidean spread of the cluster (distance of the individual IC locations from the cluster centroid divided by the number of ICs), the mean RV of the ICs in the cluster, the distance of the cluster centroid from the ROI, and the Mahalanobis distance from the median of the multivariate distribution of all cluster solutions. The last measure shows how normal, or representative for the entire distribution, the given cluster solution is. Ideally, we are looking for a solution that contains as many subjects as possible (so the final measures are representative of the group), few ICs per subject (because it is difficult to interpret several ICs per subject in an identical cortical area), a small distance from the ROI, a low spread (tight cluster around the ROI restricting it to one “functional” cortical area), a low mean RV (reflecting physiologically plausible ICs), and a low Mahalanobis distance from the median (attenuating outlier clusters). To this end, the quality measures are assigned a weight and the clustering solutions are sorted according to their summed score. The solution with the highest combined score is then taken as the final clustering solution that can be used for further analysis.

To make sure that no outlier solution is taken as the final solution, on the one hand, one can weigh the Mahalanobis distance more negatively, on the other hand, we provide plots of the locations and average scalp topographies of the five highest-ranking solutions (figure 7b). These should look very similar, which indicates that the results are stable. Depending on the ROI it might be possible to achieve a stable solution with only 100 repetitions (e.g. in the visual cortex), but deeper ROIs like the retrosplenial complex may require several thousand repetitions. If one is interested not only in one ROI but several, two options are possible: i) Optimize separately for all ROIs and create different STUDY files accordingly. A limitation of this approach is that the same IC may be present in two or more ROIs if they are too close together. ii) If one ROI is more important than the others, it might be better to only optimize for that one ROI and use this cluster solution for all subsequent analyses (Gramann et al., 2021).

## Miscellaneous functions

As a final element, the pipeline comes with a set of post-processing functionalities not pertaining to event-related EEG analysis: Different spatial filtering techniques are implemented that do not rely on blind source separation but instead make use of the additional data modalities or other information, and scripts to visualize motion and eye gaze data in an intuitive way are provided.

### Additional spatial filtering approaches

While blind source separation methods such as ICA can be beneficial in disentangling the mix of electrical sources in EEG data in general, other spatial filtering methods that make use of additional information may prove to be more powerful in circumstances where knowledge or expectations about the data are already available before the analysis. We thus provide several such options in the function *bemobil_signal_decomposition_extended*: Source Power Comodulation (SPoC; Dähne, Meinecke, et al., 2014) allows the separation of data subspaces that describe the modulation of a given target value and can, for example, be used to extract motion-related information from EEG when motion data is available, allowing either the removal or the interpretation of the data (Gehrke et al., 2019). Canonical Correlation Analysis (CCA) can be used to find common subspaces between EEG and other data, allowing the investigation of their relationships such as the interplay of EEG and functional magnetic resonance imaging (fMRI) data (Biebmann et al., 2010), or the removal of motion artifacts in EEG data (Safieddine et al., 2012). Lastly, Spatiospectral Decomposition (SSD; Nikulin et al., 2011) is a possible preprocessing step to reduce dimensionality before applying SPoC or CCA (Dähne, Nikulin, et al., 2014) but can also be used standalone to extract spatial filters that enhance specific frequencies such as the theta or alpha band in the EEG. As a final element when using the above described spatial filtering techniques, the function *bemobil_distributed_source_localization* allows the inspection of the computed spatial patterns on the source level by using previously found source locations of the ICA, a method that was originally intended to visualize the sources of brain-computer interface classifiers (Krol et al., 2018; Zander et al., 2016).

### Visualizations of motion and eye gaze data

Motion datasets can in particular be difficult to visualize without neglecting parameters that could lead to serendipitous discoveries. We have thus focused on developing multiple informative plots for both eye-tracking and motion datasets that ensure an accessible, coherent, and rapid inspection of the data. As both eye-tracking and motion data are best examined using several parameters in a single plot, we have developed plots that readily and intuitively visualize velocities and positions in space over time (figure 8).

**Figure 8.**
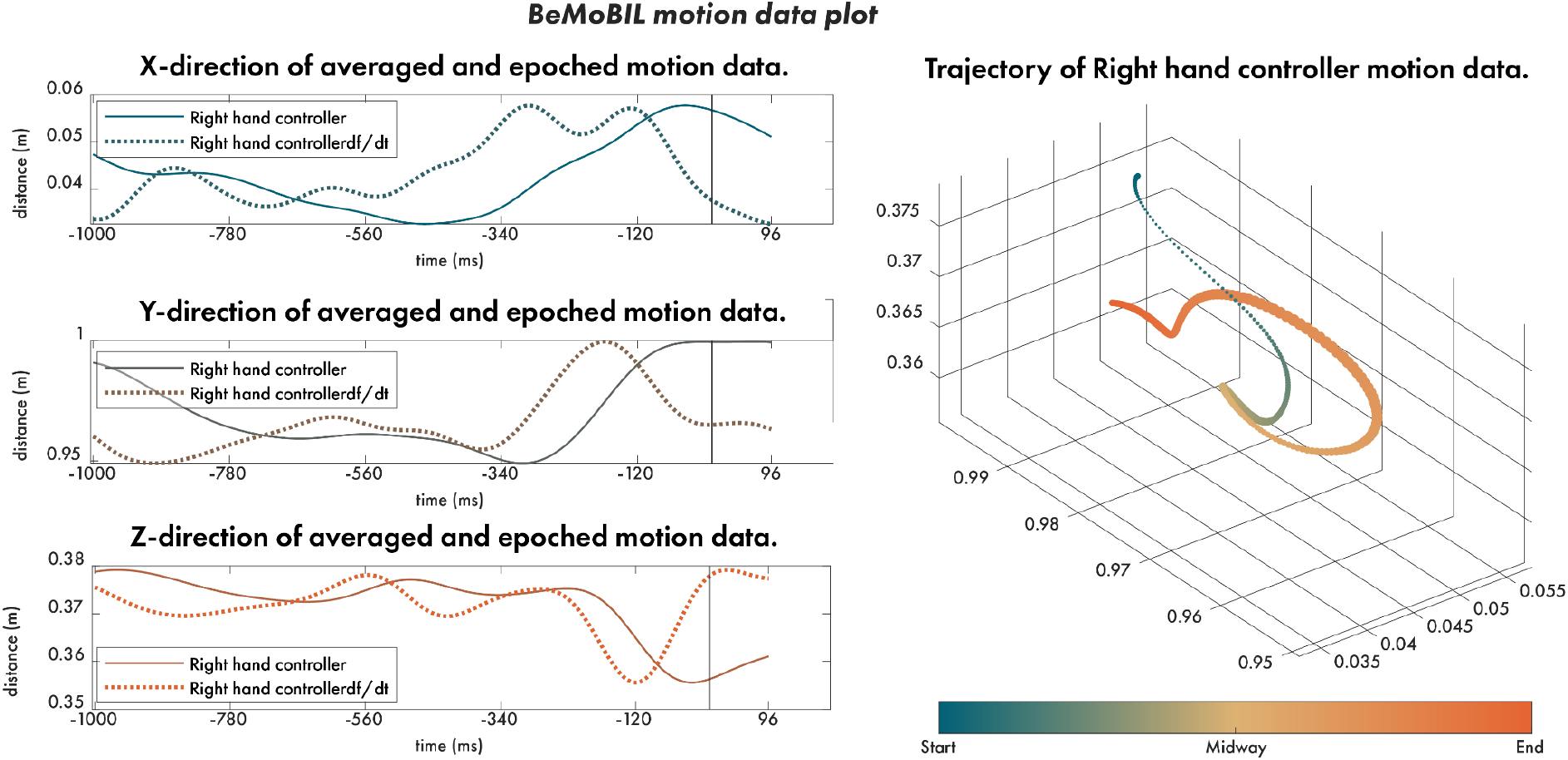
BeMoBIL motion data plot example. In this example, we visualize the controller movement in the time up to a touch of an object in a physical reaching task (data from subject 1 in the NeSitA example data that is available with the BeMoBIL Pipeline). On the left side of the plot, we visualize the synchronized XYZ coordinates separately with their velocity measures plotted on top. On the right side of the plot, we visualize the three-dimensional movement as a trajectory through space. Besides illustrating the position in space, we also show the temporal dimension using colors and the velocity of the movement using the thickness of the line as inspired by the kinematics of handwriting, supporting an intuitive and accessible reading. All the information used in the plots is available as the function output upon plotting the figure.

The pipeline provides three such functions for plotting. The first, *bemobil_plot_motion*, serves to inspect each XYZ coordinate as well as their three-dimensional trajectory. On the left side of the plot, the function divides the XYZ coordinates of the motion data into three separate plots that visualize the distance against the time as well as the velocity of the movement. The right side of the plot visualizes the three-dimensional trajectory in space without neglecting velocity or time. To visualize velocity, we have been inspired by the everyday kinematics of handwriting, i.e. strokes that are thicker relative to the rest of the line represent a slower hand movement as opposed to the thinner part of the line, representing much faster movements. To visualize the temporal dimension, we have been inspired by the techniques in plots of imaginary numbers and Riemann topology, i.e. we use a gradient of color to depict the end and beginning of the trajectory. All the information used in the plots is available as the function output upon plotting the figure.

The last two functions are dedicated to eye-tracking. Following the same principles as above, the function *bemobil_plot_trail* compares the trails of two conditions in two dimensions, using again the thickness of the trail to represent the velocity and the colors for the temporal dimension (figure 9a). As an additional plot that enables image-based analyses, we have included a heat-map function, *bemobile_plot_heatmap*, that plots the areas in which the trail spent the longest time (figure 9b). For custom colormaps, we offer the *bemobil_makecmap* which generates a gradient between given colors. The output here can be used with all our plot functions.

**Figure 9.**
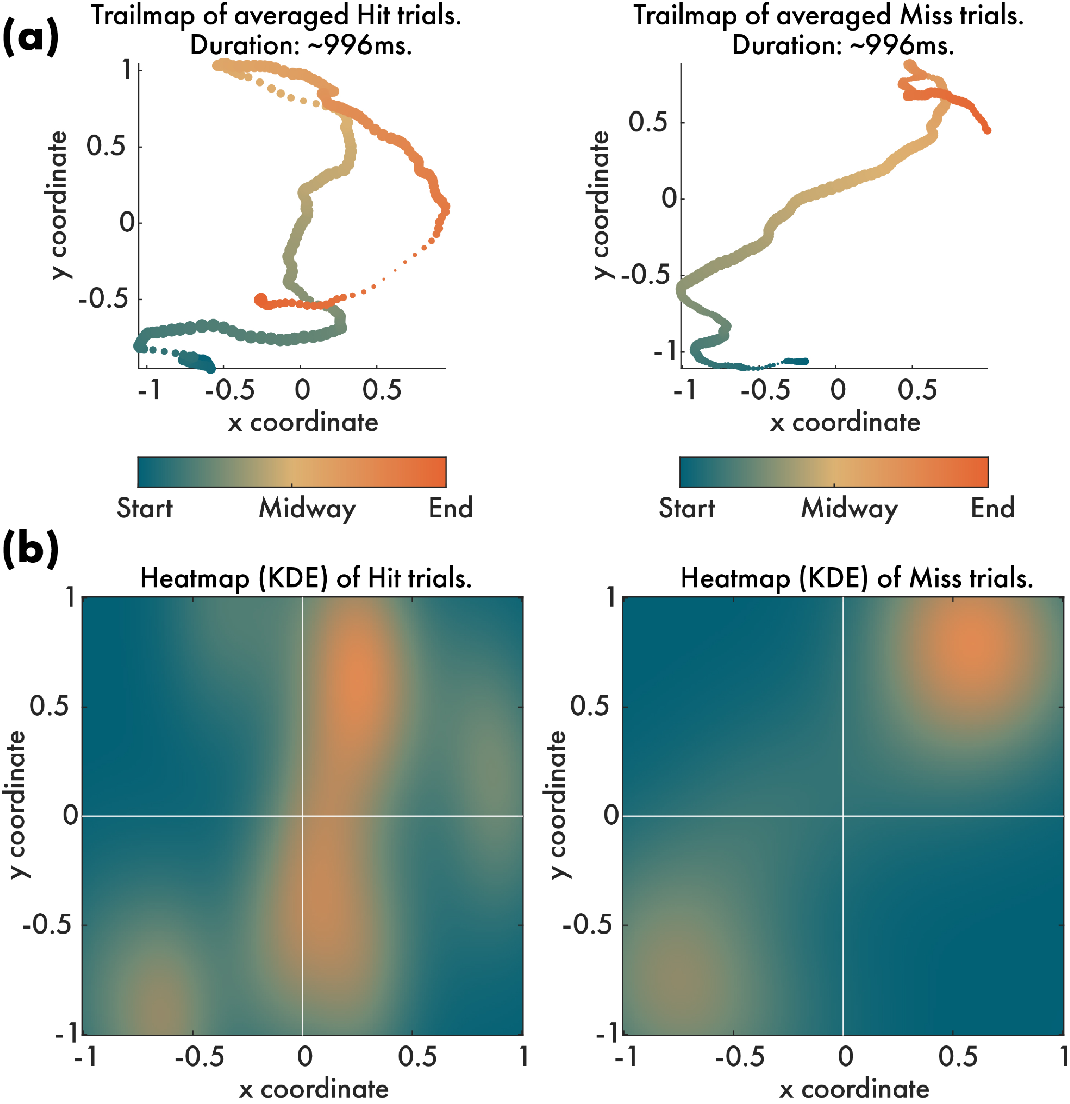
Eye tracking data plot example. (a) In this example, we compare two conditions in which the participant either hit or missed the target in a physical reaching task (data from subject 1 in the NeSitA example data that is available with the BeMoBIL Pipeline). The plot is based on the same principles as the motion data plot (see figure 8). (b) The data can readily be transferred to a heatmap kernel density estimation (KDE) plot that can be exported and further processed using image-based analyses.

## Summary

The combination of electrophysiological data and other body measures such as motion or eye tracking becomes more prevalent as a tool in neuroscientific studies investigating human brain dynamics in more ecologically valid scenarios such as the workplace (Ayaz & Dehais, 2018; Mehta & Parasuraman, 2013; Parasuraman & Rizzo, 2007; Wascher et al., 2014) or urban environments (Aspinall et al., 2015; Djebbara et al., 2019). These large multimodal datasets require special treatment during the analysis to ensure the reliability of the results. While there are powerful solutions available for individual aspects of EEG data preprocessing (e.g. the PREP pipeline (Bigdely-Shamlo et al., 2015)) or automated EEG analysis (e.g. the HAPPE pipeline (Gabard-Durnam et al., 2018)), they do not cater to the specific needs of multimodal MoBI data. For example, these pipelines do not include the synchronized import of multiple data streams, the processing of data not stemming from EEG, the treatment of EEG data from mobile experiments specifically, or additional functionalities for extracting event markers, reliable source analysis, or early-fusion analysis that can greatly benefit MoBI research.

Our proposed BeMoBIL Pipeline thus seeks to fill this gap and provides automatic, transparent, replicable, and easy-to-use data processing for multimodal datasets of human participants. It is a MATLAB pipeline based on EEGLAB (Delorme et al., 2011; Delorme & Makeig, 2004) and fieldtrip (Oostenveld et al., 2011), comprising wrappers and new functions to i) import the raw multimodal dataset to obtain BIDS-compatible shareable data and fully synchronized multimodal EEGLAB files that allow easy computations of multimodal analysis or transfer of event markers, ii) preprocess and clean EEG data, including line-noise removal, channel rejection and interpolation, and artifact rejection using ICA and epoch cleaning, iii) process motion data and extract event markers from motion, eye tracking, ECG, and IC activity, iv) robust clustering of ICs for group-level source analysis, v) allow early-fusion analysis of EEG and other data combined, and vi) visualize motion and eye gaze data intuitively. This set of features helps to reduce experimenter bias when analyzing data, allows the easy replication of data processing, and lowers the threshold of entry into EEG data analysis for researchers from other fields. Importantly, while the pipeline is designed around the requirements of MoBI studies, it can be applied to datasets from stationary studies with minimal changes.

The pipeline can be used with a small set of wrapper functions and a configuration file, but the steps can also be called individually to allow a modular setup of self-defined pipelines. For reporting the methods, all steps are documented within the EEG data structure itself, allowing precise descriptions of all data processing such as exact filter specifications, removed channels, or the amount of data removed by AMICA, to simplify replications or similar investigations in future studies. Furthermore, once set up, the entire processing from the raw files to the cleaned datasets and even the final study with clusters of interest can easily be reproduced using the pipeline scripts. This will yield identical final datasets and results though minor deviations due to suboptimal parameter selection in the configuration files (e.g., number of iterations for specific processing steps) are possible. The pipeline is documented in depth in a wiki section on the public GitHub repository, available online at https://github.com/BeMoBIL/bemobil-pipeline/wiki. Here, a comprehensive user guide on all steps including the installation can be found alongside explanations of the chosen default values and other practical considerations when running the functions.

### Limitations and conclusions

A major limitation of largely automated data processing is that researchers have no insight into the actual data and its subtleties. This can lead to overlooking processing errors, such as mishandling artifacts or unrealistic results. Therefore, we strongly believe that the visualization and close inspection of the data is an essential element of electrophysiological data analysis. To this end, analytics plots of the data during and after processing steps, as well as figures for the processing itself are created throughout the pipeline in order to keep the experimenters informed about the analysis. A comprehensive guide on the interpretation of the analytics plots is available on the wiki. Additionally, Even though the pipeline is as flexible as possible and has been extensively tested, it is in the nature of MoBI experiments that they sometimes raise unprecedented analysis issues that are difficult to predict and may require a customized pipeline. This problem can be addressed by taking modular elements of the pipeline and setting up custom analyses or adding entirely new elements as appropriate. Since the BeMoBIL Pipeline is an open source project, new analysis approaches from the community as well as contributions to the code repository are strongly encouraged. Our aim is to continuously provide the best possible pipeline that takes into account algorithmic advances or new insights while providing a stable and reliable analysis that can be easily used.

Taken together, our two main goals were to make the processing and analysis of (mobile) EEG data more reliable and independent of the researcher and to open up EEG research to other fields as a method to answer their own research questions. We provide a flexible and powerful open-source toolset for multimodal data processing and cleaning that paves the way for fast and reliable research using mobile EEG and complementary body measures.

## Supporting information

Supplementary results comparing time-cleaning types

## Acknowledgments

We gratefully acknowledge all users and testers of earlier versions of the pipeline in recent years. They provided invaluable feedback that allowed the pipeline to grow to its current state. This work has been supported by the DFG (grant number GR2627/8-1) and USAF (grant number ONR 10024807).

## Data and Code Availability Statement

The source code of the BeMoBIL Pipeline is available at https://github.com/BeMoBIL/bemobil-pipeline. The data and scripts that were used in the examples are available for download as described in the repository.

## Ethics Statement

The experimental procedures of the example data were approved by the local ethics committee (Technische Universität Berlin, Germany), the research was performed in accordance with the ethics guidelines of the Declaration of Helsinki, and all participants signed a written informed consent prior to participation.

## References

Acar, Z. A., & Makeig, S. (2010). Neuroelectromagnetic Forward Head Modeling Toolbox. Journal of Neuroscience Methods, 190(2), 258–270.

Aspinall, P., Mavros, P., Coyne, R., & Roe, J. (2015). The urban brain: analysing outdoor physical activity with mobile EEG. British Journal of Sports Medicine, 49(4), 272–276.

Ayaz, H., & Dehais,F. (Eds.). (2018). Neuroergonomics -The Brain at Work and in Everyday Life (1st ed., pp. 59–63). Elsevier Academic Press.

Bell, A. J., & Sejnowski, T. J. (1995). An information-maximization approach to blind separation and blind deconvolution. Neural Computation, 7(6), 1129–1159.

Biebmann, F., Meinecke, F. C., Gretton, A., Rauch, A., Rainer, G., Logothetis, N. K., & Müller, K. R. (2010). Temporal kernel CCA and its application in multimodal neuronal data analysis. Machine Learning, 79(1-2), 5–27.

Bigdely-Shamlo, N., Mullen, T., Kothe, C., Su, K.-M., & Robbins, K. A. (2015). The PREP pipeline: standardized preprocessing for large-scale EEG analysis. Frontiers in Neuroinformatics, 9, 16.

Büchel, D., Sandbakk, Ø., & Baumeister, J. (2021). Exploring intensity-dependent modulations in EEG resting-state network efficiency induced by exercise. European Journal of Applied Physiology, 121(9), 2423–2435.

Bulling, A., Ward, J. A., Gellersen, H., & Tröster, G. (2011). Eye movement analysis for activity recognition using electrooculography. IEEE Transactions on Pattern Analysis and Machine Intelligence, 33(4), 741–753.

Castermans, T., Duvinage, M., Cheron, G., & Dutoit, T. (2014). About the cortical origin of the low-delta and high-gamma rhythms observed in EEG signals during treadmill walking. Neuroscience Letters, 561, 166–170.

Chaumon, M., Bishop, D. V. M., & Busch, N. A. (2015). A practical guide to the selection of independent components of the electroencephalogram for artifact correction. Journal of Neuroscience Methods, 250, 47–63.

Cohen, M. X. (2017). Rigor and replication in time-frequency analyses of cognitive electrophysiology data. International Journal of Psychophysiology: Official Journal of the International Organization of Psychophysiology, 111, 80–87.

da Cruz, J. R., Chicherov, V., Herzog, M. H., & Figueiredo, P. (2018). An automatic pre-processing pipeline for EEG analysis (APP) based on robust statistics. Clinical Neurophysiology: Official Journal of the International Federation of Clinical Neurophysiology, 129(7), 1427–1437.

Dähne, S., Meinecke, F. C., Haufe, S., Höhne, J., Tangermann, M., Müller, K. R., & Nikulin, V. V. (2014). SPoC: A novel framework for relating the amplitude of neuronal oscillations to behaviorally relevant parameters. NeuroImage, 86, 111–122.

Dähne, S., Nikulin, V. V., Ramírez, D., Schreier, P. J., Müller, K. R., & Haufe, S. (2014). Finding brain oscillations with power dependencies in neuroimaging data. NeuroImage, 96, 334–348.

de Cheveigné, A. (2020). ZapLine: A simple and effective method to remove power line artifacts. NeuroImage, 207, 116356.

Delliaux, S., Delaforge, A., Deharo, J.-C., & Chaumet, G. (2019). Mental Workload Alters Heart Rate Variability, Lowering Non-linear Dynamics. Frontiers in Physiology, 10, 565.

Delorme, A., & Makeig, S. (2004). EEGLAB: An open source toolbox for analysis of single-trial EEG dynamics including independent component analysis. Journal of Neuroscience Methods, 134(1), 9–21.

Delorme, A., Mullen, T., Kothe, C., Akalin Acar, Z., Bigdely-Shamlo, N., Vankov, A., & Makeig, S. (2011). EEGLAB, SIFT, NFT, BCILAB, and ERICA: new tools for advanced EEG processing. Computational Intelligence and Neuroscience, 2011, 130714.

Delorme, A., Palmer, J., Onton, J., Oostenveld, R., & Makeig, S. (2012). Independent EEG sources are dipolar. PloS One, 7(2), e30135.

De Sanctis, P., Malcolm, B. R., Mabie, P. C., Francisco, A. A., Mowrey, W. B., Joshi, S., Molholm, S., & Foxe, J. J. (2020). Mobile Brain/Body Imaging of cognitive-motor impairment in multiple sclerosis: Deriving EEG-based neuro-markers during a dual-task walking study. Clinical Neurophysiology: Official Journal of the International Federation of Clinical Neurophysiology, 131(5), 1119–1128.

Djebbara, Z., Fich, L. B., Petrini, L., & Gramann, K. (2019). Sensory-motor brain dynamics reflect architectural affordances. Proceedings of the National Academy of Sciences, 1–31.

Evans, A. C., Collins, D. L., & Mills, S. R. (1993). 3D statistical neuroanatomical models from 305 MRI volumes. 1993 IEEE. https://ieeexplore.ieee.org/abstract/document/373602/

Gabard-Durnam, L. J., Leal, A. S. M., Wilkinson, C. L., & Levin, A. R. (2018). The harvard automated processing pipeline for electroencephalography (HAPPE): Standardized processing software for developmental and high-artifact data. Frontiers in Neuroscience, 12. https://doi.org/10.3389/fnins.2018.00097

Gehrke, L., Guerdan, L., & Gramann, K. (2019). Extracting Motion-Related Subspaces from EEG in Mobile Brain/Body Imaging Studies using Source Power Comodulation. International IEEE/EMBS Conference on Neural Engineering, NER, 2019-March, 344–347.

Gehrke, L., Lopes, P., Klug, M., Akman, S., & Gramann, K. (2022). Neural sources of prediction errors detect unrealistic VR interactions. Journal of Neural Engineering, 19(3), 036002.

Gorgolewski, K. J., Auer, T., Calhoun, V. D., Craddock, R. C., Das, S., Duff, E. P., Flandin, G., Ghosh, S. S., Glatard, T., Halchenko, Y. O., Handwerker, D. A., Hanke, M., Keator, D., Li, X., Michael, Z., Maumet, C., Nichols, B. N., Nichols, T. E., Pellman, J., … Poldrack, R. A. (2016). The brain imaging data structure, a format for organizing and describing outputs of neuroimaging experiments. Scientific Data, 3(1), 1–9.

Gramann, K., Ferris, D. P., Gwin, J., & Makeig, S. (2014). Imaging natural cognition in action. International Journal of Psychophysiology: Official Journal of the International Organization of Psychophysiology, 91(1), 22–29.

Gramann, K., Gwin, J. T., Ferris, D. P., Oie, K., Jung, T. P., Lin, C. T., Liao, L. D., & Makeig, S. (2011). Cognition in action: Imaging brain/body dynamics in mobile humans. Reviews in the Neurosciences, 22(6), 593–608.

Gramann, K., Hohlefeld, F. U., Gehrke, L., & Klug, M. (2021). Human cortical dynamics during full-body heading changes. Scientific Reports, 11(1), 18186.

Gwin, J. T., Gramann, K., Makeig, S., & Ferris, D. P. (2010). Removal of movement artifact from high-density EEG recorded during walking and running. Journal of Neurophysiology, 103(6), 3526–3534.

Harmening, N., Klug, M., Gramann, K., & Miklody, D. (2022). HArtMuT -Modeling eye and muscle contributors in neuroelectric imaging. In bioRxiv (p. 2022.08.19.504507). https://doi.org/10.1101/2022.08.19.504507

Hollman, J. H., McDade, E. M., & Petersen, R. C. (2011). Normative spatiotemporal gait parameters in older adults. Gait & Posture, 34(1), 111–118.

Hyvärinen, A., Karhunen, J., & Oja, E. (2001). Independent Component Analysis. John Wiley & Sons.

Hyvärinen, A., & Oja, E. (1997). A fast fixed-point algorithm for independent component analysis. Neural Computation, 9, 1483–1492.

Jacobsen, N. S. J., Blum, S., Witt, K., & Debener, S. (2021). A walk in the park? Characterizing gait-related artifacts in mobile EEG recordings. The European Journal of Neuroscience, 54(12), 8421–8440.

Jia, Y., & Tyler, C. W. (2019). Measurement of saccadic eye movements by electrooculography for simultaneous EEG recording. Behavior Research Methods, 51(5), 2139–2151.

Jungnickel, E., Gehrke, L., Klug, M., & Gramann, K. (2019). Chapter 10 -MoBI—Mobile Brain/Body Imaging. In H. Ayaz & F. Dehais (Eds.), Neuroergonomics (pp. 59–63). Academic Press.

Jungnickel, E., & Gramann, K. (2016). Mobile Brain/Body Imaging (MoBI) of Physical Interaction with Dynamically Moving Objects. Frontiers in Human Neuroscience, 10(June), 306.

Kappenman, E. S., & Keil, A. (2017). Introduction to the special issue on recentering science: Replication, robustness, and reproducibility in psychophysiology. Psychophysiology, 54(1), 3–5.

Kim, H.-G., Cheon, E.-J., Bai, D.-S., Lee, Y. H., & Koo, B.-H. (2018). Stress and Heart Rate Variability: A Meta-Analysis and Review of the Literature. Psychiatry Investigation, 15(3), 235–245.

Klug, M., Berg, T., & Gramann, K. (2022). No need for extensive artifact rejection for ICA -A multi-study evaluation on stationary and mobile EEG datasets. In bioRxiv (p. 2022.09.13.507772). https://doi.org/10.1101/2022.09.13.507772

Klug, M., & Gramann, K. (2021). Identifying key factors for improving ICA-based decomposition of EEG data in mobile and stationary experiments. The European Journal of Neuroscience, 54(12), 8406–8420.

Klug, M., & Kloosterman, N. A. (2022). Zapline-plus: A Zapline extension for automatic and adaptive removal of frequency-specific noise artifacts in M/EEG. Human Brain Mapping, 43(9), 2743–2758.

Krol, L., Mousavi, M., De Sa, V., & Zander, T. (2018). Towards Classifier Visualisation in 3D Source Space. 2018 IEEE International Conference on Systems, Man, and Cybernetics (SMC), 71–76.

Larson, M. J., & Moser, J. S. (2017). Rigor and replication: Toward improved best practices in human electrophysiology research. International Journal of Psychophysiology: Official Journal of the International Organization of Psychophysiology, 111, 1–4.

Leutheuser, H., Gabsteiger, F., Hebenstreit, F., Reis, P., Lochmann, M., & Eskofier, B. (2013). Comparison of the AMICA and the InfoMax algorithm for the reduction of electromyogenic artifacts in EEG data. Proceedings of the Annual International Conference of the IEEE Engineering in Medicine and Biology Society, EMBS, 6804–6807.

Lins, O. G., Picton, T. W., Berg, P., & Scherg, M. (1993). Ocular artifacts in EEG and event-related potentials. I: Scalp topography. Brain Topography, 6(1), 51–63.

Makeig, S., Gramann, K., Jung, T.-P., Sejnowski, T. J., & Poizner, H. (2009). Linking brain, mind and behavior. International Journal of Psychophysiology: Official Journal of the International Organization of Psychophysiology, 73(2), 95–100.

Maki, B. E. (1997). Gait changes in older adults: predictors of falls or indicators of fear. Journal of the American Geriatrics Society, 45(3), 313–320.

Malcolm, B. R., Foxe, J. J., Butler, J. S., & De Sanctis, P. (2015). The aging brain shows less flexible reallocation of cognitive resources during dual-task walking: A mobile brain/body imaging (MoBI) study. NeuroImage, 117, 230–242.

Mehta, R. K., & Parasuraman, R. (2013). Neuroergonomics: a review of applications to physical and cognitive work. Frontiers in Human Neuroscience, 7(December), 1–10.

Mitz, A. R., Chacko, R. V., Putnam, P. T., Rudebeck, P. H., & Murray, E. A. (2017). Using pupil size and heart rate to infer affective states during behavioral neurophysiology and neuropsychology experiments. Journal of Neuroscience Methods, 279. https://doi.org/10.1016/j.jneumeth.2017.01.004

Miyakoshi, M., Schmitt, L. M., Erickson, C. A., Sweeney, J. A., & Pedapati, E. V. (2021). Can We Push the “Quasi-Perfect Artifact Rejection” Even Closer to Perfection? Frontiers in Neuroinformatics, 14(January), 1–5.

Nenna, F., Do, C. T., Protzak, J., & Gramann, K. (2021). Alteration of brain dynamics during dual-task overground walking. The European Journal of Neuroscience, 54(12), 8158–8174.

Nikulin, V. V., Nolte, G., & Curio, G. (2011). A novel method for reliable and fast extraction of neuronal EEG / MEG oscillations on the basis of spatio-spectral decomposition. NeuroImage, 55(4), 1528–1535.

Ojeda, A., Bigdely-Shamlo, N., & Makeig, S. (2014). MoBILAB: an open source toolbox for analysis and visualization of mobile brain/body imaging data. Frontiers in Human Neuroscience, 8(March), 121.

Oostenveld, R., Fries, P., Maris, E., & Schoffelen, J.-M. (2011). FieldTrip: Open source software for advanced analysis of MEG, EEG, and invasive electrophysiological data. Computational Intelligence and Neuroscience, 2011, 156869.

Open Science Collaboration. (2015). Estimating the reproducibility of psychological science. Science, 349(6251). https://doi.org/10.1126/science.aac4716

Palmer, J. A., Kreutz-delgado, K., & Makeig, S. (2011). AMICA : An Adaptive Mixture of Independent Component Analyzers with Shared Components. 1–15.

Pan, J., & Tompkins, W. J. (1985). A real-time QRS detection algorithm. IEEE Transactions on Bio-Medical Engineering, 32(3), 230–236.

Parasuraman, R., & Rizzo, M. (Eds.). (2007). Neuroergonomics -The brain at work. Oxford University Press.

Pedroni, A., Bahreini, A., & Langer, N. (2019). Automagic: Standardized preprocessing of big EEG data. NeuroImage, 200(December 2018), 460–473.

Pernet, C. R., Appelhoff, S., Gorgolewski, K. J., Flandin, G., Phillips, C., Delorme, A., & Oostenveld, R. (2019). EEG-BIDS, an extension to the brain imaging data structure for electroencephalography. Scientific Data, 6(1), 1–5.

Pernet, C. R., Martinez-Cancino, R., Truong, D., Makeig, S., & Delorme, A. (2021). From BIDS-Formatted EEG Data to Sensor-Space Group Results: A Fully Reproducible Workflow With EEGLAB and LIMO EEG. Frontiers in Neuroscience, 14(January), 1–7.

Perry, J., & Burnfield, J. (2010). Gait analysis. Normal and pathological function (2nd ed.). SLACK Inc.

Pion-Tonachini, L., Kreutz-Delgado, K., & Makeig, S. (2019). ICLabel: An automated electroencephalographic independent component classifier, dataset, and website. NeuroImage, 198(May), 181–197.

Protzak, J., & Gramann, K. (2018). Investigating established EEG parameter during real-world driving. Frontiers in Psychology, 9(NOV), 1–11.

Protzak, J., & Gramann, K. (2021). EEG beta-modulations reflect age-specific motor resource allocation during dual-task walking. Scientific Reports, 11(1), 16110.

Protzak, J., Wiczorek, R., & Gramann, K. (2021). Peripheral visual perception during natural overground dual-task walking in older and younger adults. Neurobiology of Aging, 98, 146–159.

Reiser, J. E., Wascher, E., & Arnau, S. (2019). Recording mobile EEG in an outdoor environment reveals cognitive-motor interference dependent on movement complexity. Scientific Reports, 9(1). https://doi.org/10.1038/s41598-019-49503-4

Reiser, J. E., Wascher, E., Rinkenauer, G., & Arnau, S. (2021). Cognitive-motor interference in the wild: Assessing the effects of movement complexity on task switching using mobile EEG. The European Journal of Neuroscience, 54(12), 8175–8195.

Richer, N., Downey, R. J., Hairston, W. D., Ferris, D. P., & Nordin, A. D. (2020). Motion and Muscle Artifact Removal Validation Using an Electrical Head Phantom, Robotic Motion Platform, and Dual Layer Mobile EEG. IEEE Transactions on Neural Systems and Rehabilitation Engineering: A Publication of the IEEE Engineering in Medicine and Biology Society, 28(8), 1825–1835.

Robbins, K. A., Touryan, J., Mullen, T., Kothe, C., & Bigdely-Shamlo, N. (2020). How Sensitive Are EEG Results to Preprocessing Methods: A Benchmarking Study. IEEE Transactions on Neural Systems and Rehabilitation Engineering: A Publication of the IEEE Engineering in Medicine and Biology Society, 28(5), 1081–1090.

Rodrigues, J., Weiß, M., Hewig, J., & Allen, J. J. B. (2021). EPOS: EEG Processing Open-Source Scripts. Frontiers in Neuroscience, 15, 660449.

Rudisch, J., Jöllenbeck, T., Vogt, L., Cordes, T., Klotzbier, T. J., Vogel, O., & Wollesen, B. (2021). Agreement and consistency of five different clinical gait analysis systems in the assessment of spatiotemporal gait parameters. Gait & Posture, 85, 55–64.

Safieddine, D., Kachenoura, A., Albera, L., Birot, G., Karfoul, A., Pasnicu, A., Biraben, A., Wendling, F., Senhadji, L., & Merlet, I. (2012). Removal of muscle artifact from EEG data: Comparison between stochastic (ICA and CCA) and deterministic (EMD and wavelet-based) approaches. EURASIP Journal on Advances in Signal Processing, 2012(1). https://doi.org/10.1186/1687-6180-2012-127

Scott, D., McLaughlin, P., Nicholson, G. C., Ebeling, P. R., Stuart, A. L., Kay, D., & Sanders, K. M. (2015). Changes in gait performance over several years are associated with recurrent falls status in community-dwelling older women at high risk of fracture. Age and Ageing, 44(2), 287–293.

Sedghamiz, H. (2014). Matlab Implementation of Pan Tompkins ECG QRS detector. ResearchGate. https://www.researchgate.net/publication/313673153_Matlab_Implementation_of_Pan_Tompkins_ECG_QRS_detector

Seeber, M., Scherer, R., Wagner, J., Solis-Escalante, T., & Müller-Putz, G. R. (2015). High and low gamma EEG oscillations in central sensorimotor areas are conversely modulated during the human gait cycle. NeuroImage, 112, 318–326.

Short, M. R., Damiano, D. L., Kim, Y., & Bulea, T. C. (2020). Children With Unilateral Cerebral Palsy Utilize More Cortical Resources for Similar Motor Output During Treadmill Gait. Frontiers in Human Neuroscience, 14, 36.

Verghese, J., Holtzer, R., Lipton, R. B., & Wang, C. (2009). Quantitative gait markers and incident fall risk in older adults. The Journals of Gerontology. Series A, Biological Sciences and Medical Sciences, 64(8), 896–901.

Wagner, J., Makeig, S., Gola, M., Neuper, C., & Muller-Putz, G. (2016). Distinct Band Oscillatory Networks Subserving Motor and Cognitive Control during Gait Adaptation. Journal of Neuroscience, 36(7), 2212–2226.

Wagner, J., Solis-Escalante, T., Grieshofer, P., Neuper, C., Müller-Putz, G., & Scherer, R. (2012). Level of participation in robotic-assisted treadmill walking modulates midline sensorimotor EEG rhythms in able-bodied subjects. NeuroImage, 63(3), 1203–1211.

Wascher, E., Heppner, H., & Hoffmann, S. (2014). Towards the measurement of event-related EEG activity in real-life working environments. International Journal of Psychophysiology: Official Journal of the International Organization of Psychophysiology, 91(1), 3–9.

Widmann, A., Schröger, E., & Maess, B. (2015). Digital filter design for electrophysiological data – a practical approach. Journal of Neuroscience Methods, 250, 34–46.

Wunderlich, A., & Gramann, K. (2021). Eye movement-related brain potentials during assisted navigation in real-world environments. The European Journal of Neuroscience, 54(12), 8336–8354.

Zakeri, Z., Assecondi, S., Bagshaw, A. P., & Arvanitis, T. N. (2014). Influence of Signal Preprocessing on ICA-Based EEG Decomposition. IFMBE Proceedings, 41, 563–566.

Zander, T. O., Krol, L. R., Birbaumer, N. P., & Gramann, K. (2016). Neuroadaptive technology enables implicit cursor control based on medial prefrontal cortex activity. Proceedings of the National Academy of Sciences, 113(52), 14898–14903.

