## Supplementary results comparing time-cleaning types for "The BeMoBIL Pipeline for automated analyses of multimodal mobile brain and body imaging data"

**Comparison of the BeMoBIL time-cleaning with the automatic AMICA sample rejection**

We decomposed 8 studies with AMICA and investigated the decomposition quality. We compared no time cleaning with the two different methods and another using both methods in succession. The data and processing are otherwise identical to that in Klug et al. (2022). For the BeMoBIL time cleaning, we used a threshold of 5%, resulting in around 6% of data removed due to added buffering. For the AMICA autoclean, we used 5 iterations and 3 standard deviations as parameters.

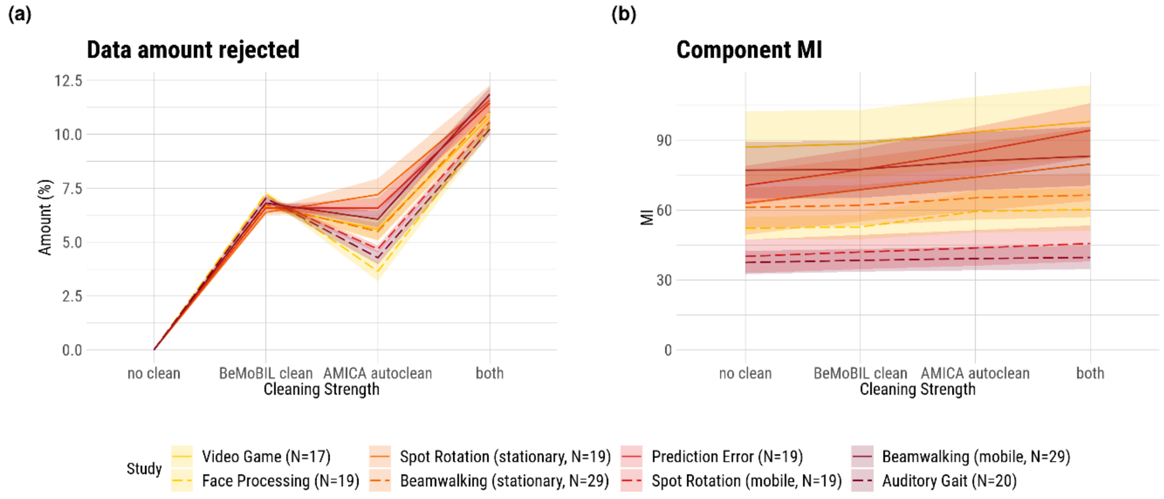


*1: Results for the data rejection amount and component mutual information (MI). Shaded areas depict the standard error of the mean (SE). "No clean" refers to no sample rejection being applied when computing AMICA. The colors denote the movement intensities: yellow - low, orange - low-to-medium, red - medium-to-high, violet - high.*

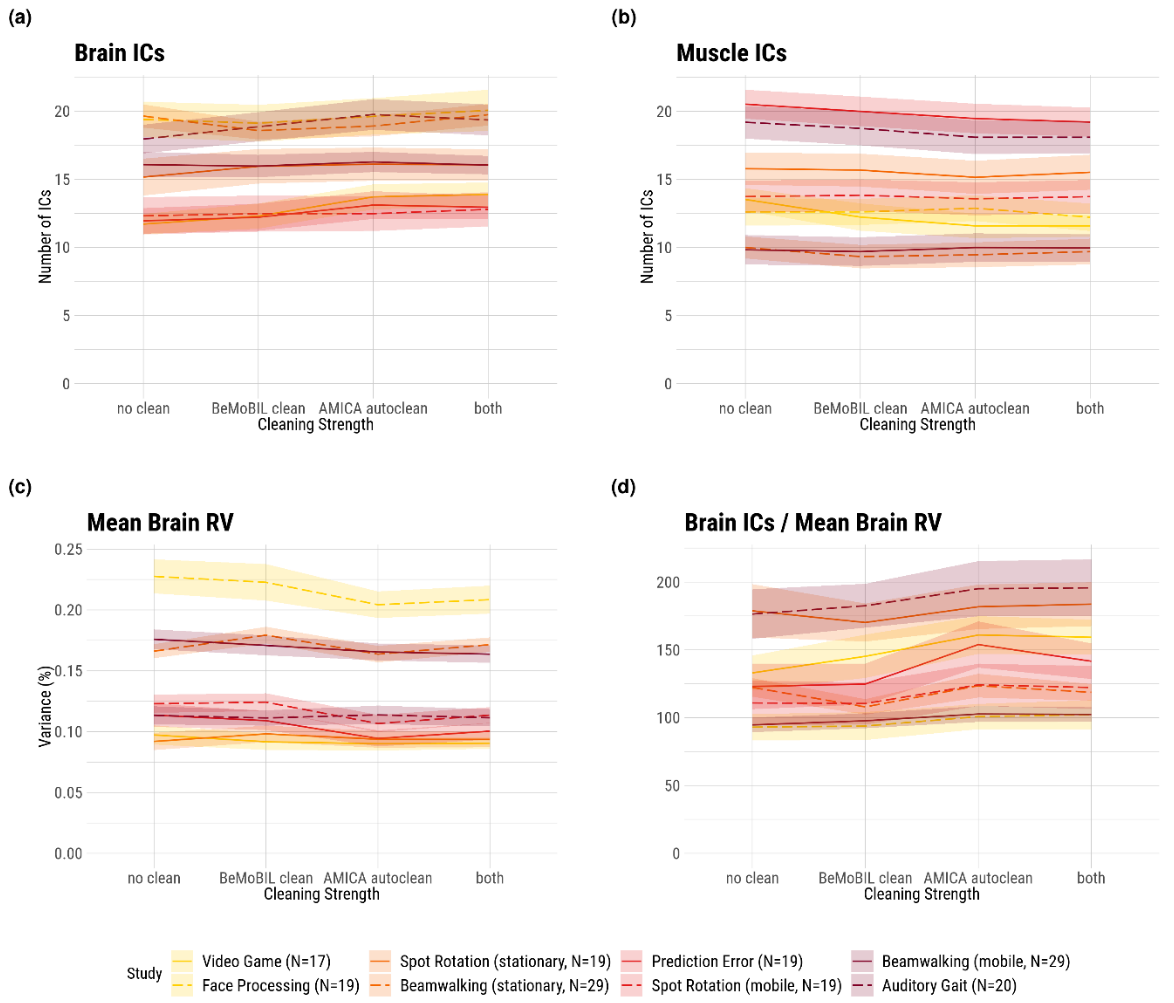


*2: Results for the ICLabel classification and residual variance (RV). Shaded areas depict the standard error of the mean (SE). "No clean" refers to no sample rejection being applied when computing AMICA. The colors denote the movement intensities: yellow - low, orange - low-to-medium, red - medium-to-high, violet - high.*

The results indicate no relevant difference between the two decomposition methods. On average, a similar amount of data was rejected, and the change in brain ICs and RV was mostly within the SE range. The small trend towards a better decomposition in the automatic AMICA sample rejection resulted in this method being chosen for the BeMoBIL Pipeline.

### References


Klug, M., Berg, T., & Gramann, K. (2022). No need for extensive artifact rejection for ICA - A multi-study evaluation on stationary and mobile EEG datasets. In *bioRxiv* (p. 2022.09.13.507772). https://doi.org/10.1101/2022.09.13.507772
